# LORE receptor homomerization is required for 3-hydroxydecanoic acid-induced immune signaling and determines the natural variation of immunosensitivity within the Arabidopsis genus

**DOI:** 10.1101/2021.09.27.461997

**Authors:** Sabine Eschrig, Milena Schäffer, Lin-Jie Shu, Tina Illig, Sonja Eibel, Atiara Fernandez, Stefanie Ranf

## Abstract

- The S-domain-type receptor-like kinase (SD-RLK) LIPOOLIGOSACCHARIDE-SPECIFIC REDUCED ELICITATION (LORE) from *Arabidopsis thaliana* is a pattern recognition receptor that senses medium-chain 3-hydroxy fatty acids, such as 3-hydroxydecanoic acid (3-OH-C10:0), to activate pattern-triggered immunity. Here, we show that LORE homomerization is required to activate 3-OH-C10:0-induced immune signaling.
- Fluorescence lifetime imaging in *Nicotiana benthamiana* demonstrated that *At*LORE homomerizes via the extracellular and transmembrane domains. Co-expression of *At*LORE truncations lacking the intracellular domain exerts a dominant negative effect on *At*LORE signaling in both *N. benthamiana* and *A. thaliana*, highlighting that homomerization is essential for signaling.
- Screening for 3-OH-C10:0-induced reactive oxygen species production revealed natural variation within the Arabidopsis genus. *Arabidopsis lyrata* and *Arabidopsis halleri* do not respond to 3-OH-C10:0, although both possess a putative LORE orthologue. Both LORE orthologues have defective extracellular domains that bind 3-OH-C10:0 to a similar level but lack the ability to homomerize. Thus, ligand binding is independent of LORE homomerization. Analysis of *At*LORE and *Alyr*LORE chimera suggests that the loss of *Alyr*LORE homomerization is caused by several amino acid polymorphisms across the extracellular domain.
- Our findings shed light on the activation mechanism of LORE and the loss of 3-OH-C10:0 perception within the Arabidopsis genus.

## INTRODUCTION

Perception, processing and integration of environmental and cellular stimuli are fundamental for all living organisms. In plants, this is mainly implemented by members of the super-families of receptor-like kinases (RLKs) and receptor-like proteins (RLPs) (Shiu & Bleecker, 2001; Shiu & Bleecker, 2003; Hohmann *et al*., 2017; Jamieson *et al*., 2018; Dievart *et al*., 2020). They regulate various cellular processes, including growth, development, reproduction, and immunity (Shiu & Bleecker, 2001; De Smet *et al*., 2009; Li & Yang, 2016; Boutrot & Zipfel, 2017; Gou & Li, 2020). Most RLKs contain an extracellular (ECD), a single-span transmembrane (TMD) and an intracellular domain (ICD). ECDs often comprise diverse sequence motifs, including a ligand binding site, whereas ICDs contain a serine/threonine or dual-specificity protein kinase for signaling (Shiu & Bleecker, 2001; Bojar *et al*., 2014; Macho *et al*., 2015; Dievart *et al*., 2020). RLPs, lacking a kinase domain, or RLKs with pseudo-kinases (Castells & Casacuberta, 2007; Paul & Srinivasan, 2020) require signaling competent partners (Jamieson *et al*., 2018). RLKs/RLPs are classified according to their ECD architecture, e.g. as leucine-rich-repeat (LRR), L-type lectin (Lec), S-domain (SD) or lysin-motif (LysM) type (Shiu & Bleecker, 2001; Shiu & Bleecker, 2003; Dievart *et al*., 2020).

A common feature of RLK/RLPs is their assembly into higher-order receptor complexes. They can act as receptors, co-receptors, scaffolds or positive/negative regulators to orchestrate and fine-tune signaling (Ma, X *et al*., 2016; Burkart & Stahl, 2017; Wan *et al*., 2019; Gou & Li, 2020). To date, we have gained a detailed molecular understanding of activation mechanisms and the role of complex formation for only a few RLKs. One prototypical example is the LRR-RLK FLAGELLIN SENSING 2 (*At*FLS2), a pattern recognition receptor (PRR) in plant immunity. *At*FLS2 associates with LRR-co-receptors of the SOMATIC EMBRYOGENESIS RECEPTOR-LIKE KINASE (*At*SERK) family, particularly BRI1-ASSOCIATED RECEPTOR KINASE 1 (*At*BAK1/SERK3) (Gomez-Gomez & Boller, 2000; Chinchilla *et al*., 2006; Chinchilla *et al*., 2007; Roux *et al*., 2011) upon sensing the bacterial flagellin epitope flg22 (Boller & Felix, 2009; Robatzek & Wirthmueller, 2013; Sun, Y *et al*., 2013; Couto & Zipfel, 2016). In *Arabidopsis*, the multicomponent PRR complex sensing fungal chitin oligomers comprises three LysM-RLKs (LYK), *At*LYK4, *At*LYK5 and the CHITIN ELICITOR RECEPTOR KINASE 1 (*At*CERK1/*At*LYK1) (Miya *et al*., 2007; Cao *et al*., 2014; Xue *et al*., 2019; Gong *et al*., 2020). Upon chitin binding, *At*LYK5 hetero-dimerizes with *At*CERK1, triggering *At*CERK1 homomerization and the formation of a sandwich-like receptor complex (Couto & Zipfel, 2016; Gong *et al*., 2020).

While in the previous examples ligand-induced hetero-dimerization with co-receptors activates signaling, the mode of action is different for homomerizing S-LOCUS RECEPTOR KINASES (SRKs) from *Brassica rapa*. *Bra*SRKs belong to the family of SD-RLKs (alias G- or B-type lectin RLKs) with about 40 members in *A. thaliana* and more than 100 in rice (Shiu & Bleecker, 2001; Shiu *et al*., 2004; Vaid *et al*., 2012; Xing *et al*., 2013; Teixeira *et al*., 2018). Stigma-expressed *Bra*SRKs and their pollen-secreted ligands, the S-LOCUS CYSTEIN-RICH PEPTIDES (SCRs), mediate self-incompatibility (SI) and avoid inbreeding through self-pollination to maintain genetic variability (Ivanov *et al*., 2010; Jany *et al*., 2019). The ECD of SD-RLKs typically comprises two G-type lectin-like, an epidermal growth factor (EGF)-like and a plasminogen-apple-nematode (PAN) domain (Shiu & Bleecker, 2001; Xing *et al*., 2013; Dievart *et al*., 2020). Apart from SRKs, only a few SD-RLKs have been described in more detail. They are involved in plant-pathogen interactions, symbiosis or abiotic stress responses (Walker & Zhang, 1990; Tobias *et al*., 1992; Tobias & Nasrallah, 1996; Chen *et al*., 2006; Kim, H. S. *et al*., 2009; Kim, Ho Soo *et al*., 2009; Gilardoni *et al*., 2011; Chen *et al*., 2013; Cheng *et al*., 2013; Sun, XL *et al*., 2013; Trontin *et al*., 2014; Ranf *et al*., 2015; Zou *et al*., 2015; Fan *et al*., 2018; Schnepf *et al*., 2018; Kutschera *et al*., 2019; Labbé *et al*., 2019; Park *et al*., 2019; Jinjun *et al*., 2020; Pan *et al*., 2020; Sun *et al*., 2020; Liu *et al*., 2021; Mondal *et al*., 2021; Schellenberger *et al*., 2021; Zhou *et al*., 2021; Kato *et al*., 2022; Ma *et al*., 2022; Bao *et al*., 2023; Pi *et al*., 2023; Shrestha *et al*., 2023). Although a growing number of members of the large SD-RLK sub-family are being characterized, their ligands, receptor complex formation and downstream signaling mechanisms remain largely unknown.

In *A. thaliana*, we discovered the SD-RLK *At*LORE as PRR for bacterial medium-chain 3-hydroxylated fatty acids (3-OH-FAs) and 3-(3-hydroxyalkanoyloxy)alkanoates (HAAs), which induce pattern-triggered immunity (PTI) (Ranf *et al*., 2015; Kutschera *et al*., 2019; Schellenberger *et al*., 2021). Only 3-OH-FAs/HAAs with acyl chains comprising 8 to 12 carbon atoms activate *At*LORE signaling. 3-hydroxydecanoic acid (3-OH-C10:0) is the strongest elicitor in *A. thaliana* and binds to the *At*LORE-ECD (Kutschera *et al*., 2019; Shu *et al*., 2021). LORE possesses a strong RD-type kinase (Johnson *et al*., 1996; Ranf *et al*., 2015), which auto-phosphorylates *At*LORE upon 3-OH-C10:0 sensing and activates the receptor-like cytoplasmic kinase PBS1-LIKE 34/35/36 (*At*PBL34/35/36) (Luo *et al*., 2020), RPM1-INDUCED PROTEIN KINASE (*At*RIPK) and the LORE-ASSOCIATED PROTEIN PHOSPHATASE (LOPP) (Wang *et al*., 2023). To date, no co-receptor has been found to be involved in *At*LORE signaling and its activation mechanism remains unknown. Phylogenetic analysis revealed that *LORE* is restricted to Brassicaceae (Ranf *et al*., 2015). As the first SD-RLK in *A. thaliana* with a known ligand and thus controllable means of activation, *At*LORE is an important model for mechanistic studies of the SD-RLK family.

We show that *At*LORE homomerizes via its TMD and ECD, which is required for 3-OH-C10:0-induced immune signaling. Interestingly, we found natural variation in responsiveness to 3-OH-C10:0 in closely related Brassicaceae, which have putative LORE orthologues with high protein sequence homology. Functional analyses of these orthologues and *At*LORE chimera show that LORE from non-responsive species have defective ECDs. Non-functional LORE orthologues bind the 3-OH-C10:0 ligand but are compromised in homomerization. In contrast to ligand- and receptor-mediated homodimerization of *Bra*SRKs, ligand binding to LORE occurs independently of homomerization. We further show that analyzing cell death-like autoimmunity phenotypes and dominant negative effects are valuable tools for studying LORE complex formation. Overall, our results provide new insights into the mechanism of LORE-mediated 3-OH-C10:0 sensing in Brassicaceae.

## MATERIALS AND METHODS

### Sequence alignments and protein models

Multiple protein sequence alignments were performed with Jalview version 2.11.1.4 (Waterhouse *et al*., 2009) using the MAFFT (multiple alignment using fast Fourier transformation) algorithm (Katoh *et al*., 2019). Sequences were retrieved from Uniprot (Consortium, 2020), Phytozome (Goodstein *et al*., 2011), iTAK (Zheng *et al*., 2016) or NCBI (Supplementary Information **Table S5**). For LORE, domains were annotated according to Uniprot (O64782) and Naithani et al., 2007 (Supplementary Information **Table S1**). Protein models were retrieved from Uniprot for *At*LORE (Alphafold model AF-O64782-F1) and Ma *et al*., 2016 for *Bra*SRK9 and analyzed with ChimeraX (version 1.7.dev202308310222) (Pettersen *et al*., 2021).

### Molecular cloning

Plasmids were cloned using Golden Gate (Engler *et al*., 2009; Weber *et al*., 2011) or Gateway™ techniques (Katzen, 2007). Coding sequences (CDS) were PCR-amplified from cDNA or available vectors and ligated into a Golden Gate-adapted pUC18 vector (Ranf *et al*., 2015). PCR primers are listed in Supplementary Information **Table S2**. Conserved ATP binding sites of LORE-ICDs (Ranf *et al*., 2015; Luo *et al*., 2020) were modified by site-directed mutagenesis as described. For *Ahal*LORE, the first intron of *At*LORE (109 bp) was amplified from genomic DNA and inserted into *Ahal*LORE CDS (at codon 45) to circumvent problematic read-through in *E. coli* or *Agrobacterium tumefaciens*. *At*LORE truncations and receptor chimera with *Alyr*LORE were designed according to domain annotations from Uniprot and Naithani *et al.,* 2007 (Supplementary Information **Table S1, S3)**. CDS parts were amplified by PCR with *Bpi*I-containing adapters and re-assembled by Golden Gate techniques (Supplementary Information **Table S2-3**). For membrane-bound truncations, the *At*LORE signal peptide was integrated. Details of expression plasmids are listed in Supplementary Information **Table S4**.

### Plant material and cultivation

*N. benthamiana*, *A. thaliana* Col-0, *A. halleri* (N9852), *A. lyrata* (MN47) and *C. rubella* (N22697) were sown on standard potting soil and vermiculite (9:1). Brassicaceae seeds were stratified (darkness, 48 h, 4°C) and grown under short-day conditions (8 h light, 21°C, 60% relative humidity). *N. benthamiana* and flowering *A. thaliana* were grown under long-day conditions (16 h light, 24°C, 60% humidity). *A. thaliana* Col-0^AEQ^ overexpressing *At*LORE (CaMV35S:*LORE*, LORE-OE) was previously described (Shu *et al*., 2021). For cytosolic [Ca^2+^] measurements, seeds were surface-sterilized with chlorine gas (4 h) and grown in liquid medium (0.5× Murashige & Skoog medium including vitamins (Duchefa), 0.25% sucrose, 1 mM MES, pH 5.7) under long-day conditions.

### Transformation of N. benthamiana and A. thaliana

Agrobacterium-mediated transformation of *N. benthamiana* and *A. thaliana* was performed as described (Ranf *et al*., 2015; Shu *et al*., 2021). *Agrobacterium tumefaciens* GV3101 pMP90 carrying expression plasmids (Supplementary Information **Table S4**) were cultivated on LB agar with 30 μg/mL gentamycin, 10 μg/mL rifampicin, 50 μg/mL kanamycin or 100 μg/mL spectinomycin. For transient transformation of *N. benthamiana*, Agrobacteria carrying desired vectors or the silencing suppressor p19 were mixed in a 1:1 ratio or 1:1:1 in case of co-expression. Agrobacteria suspensions were adjusted to an OD_600_ of 0.5 or to 0.025 for GOF-ROS measurements to avoid cell death. For competition assays (**Fig 7**), the OD_600_ was adjusted to 0.025 and candidates mixed in the ratio 6:1:5 (p19:*At*LORE:competing component) (**Fig 7A-B**) or to OD 1.0 and mixed in the ratio 3:1:5 (**Fig 7C-D**). Agro-infiltrated plants were incubated for two days for Co-IPs, four days for chlorophyll fluorescence and 30-36 hours for GOF-ROS measurements. Expression of estradiol-inducible FLIM constructs was induced by infiltration of 15-20 µM β-estradiol and 0.1% Tween 20 into transformed leaf areas 24 h after Agro-infiltration. FLIM was performed 18-28 h after induction. Stable transgenic *A. thaliana* were generated by floral-dipping into Agrobacteria suspension (OD_600_=2.0, 0.5% sucrose, 0.03% Silwet L-77). Transformants were selected via FastRed fluorescence (Shimada *et al*., 2010; Ursache *et al*., 2021).

### Protein extraction

Plant material (60-70 leaf discs, Ø 4 mm) was frozen in liquid nitrogen and ground to fine powder (TissueLyzerII, Qiagen, 1 min, 30 Hz). Total protein was extracted by incubation with extraction buffer (6 µL/leaf disc, 150 mM Tris-HCl pH 7.5, 150 mM NaCl, 10% glycerol, 1% Nonidet-P40, 10 mM EDTA, 1 mM Na_2_MoO_4_, 1 mM NaF, 1 mM DTT, 1% (w/v) polyvinylpyrrolidone, 1% (v/v) protease inhibitor cocktail P9599 (Sigma-Aldrich), 1 h, 4°C). After centrifugation (18000 g, 30 min, 4°C), the supernatant was used for immunoblot or Co-IP experiments. For competition assays (**Fig S10**), proteins were extracted with the Minute™ Plant Plasma Membrane Protein Isolation Kit (Invent Biotechnologies, Inc., SM-005). Membrane pellets were resuspended in 25 μL 1xSDS-sample buffer (12 mM Tris-HCL pH 6.8, 0.4% SDS, 2% glycerol, 1% β-mercaptoethanol, 0.002% bromophenol blue).

### Co-immunoprecipitation

Protein extracts were incubated (1-2 h, 4°C) with GFP-Trap_MA (magnetic beads, ChromoTek) according to manufacturer’s instructions. Magnetic beads were washed three times with buffer (150 mM Tris-HCL pH 7.5, 150 mM NaCl, 0.5% Nonidet-P40) and re-suspended in 20 μL 1xSDS-sample buffer.

### Immunoblot

Samples were denatured (10 min, 95 °C) and separated by SDS-polyacrylamide gel electrophoresis (5% stacking gel, 10% resolving gel, 60-100 V, 1x Laemmli buffer). Proteins were blotted onto 0.2 μm Protran™ nitrocellulose (GE healthcare) or PVDF membrane (Merck) by semi-dry transfer (Bio-Rad, 1 mA/cm^2^, 1 h) with 1x transfer buffer (3.03 g/L Tris base, 14.4 g/L glycine, 20% methanol, 0.05% SDS). Membranes were blocked with 3% milk powder, protein-free blocking solution T20 (Pierce) or BlueBlock PF (SERVA) and incubated with respective primary antibodies diluted in blocking solution (1 h). After washing with TBS-T (6.06 g/L Tris base, 8.76 g/L NaCl, pH 6.7, 0.05% Tween20), membranes were incubated with secondary antibodies diluted in TBS-T (1 h). Antibodies are listed in Supplementary Information **Table S6**. After washing of membranes (3x 10 min, TBS-T), chemiluminescence was detected with a CCD camera (Fusion SL System, Vilber Lourmat GmbH) upon incubation with peroxidase substrates (SuperSignal®West Femto Maximum Sensitivity Substrate, SuperSignal®West Dura Extended Duration Substrate, Pierce).

When immunoblots were re-analyzed with a different antibody, membranes were washed with TBS-T, incubated in stripping buffer (2% SDS, 62.5 mM Tris/HCl pH 6.7, 100mM β-mercaptoethanol, 50°C, 30 min), washed with TBS-T, blocked and immuno-detected again. Membranes were stained for total protein with amido black (1% (w/v) amido black, 25% (v/v) isopropyl, 10% (v/v) acetic acid) or Coomassie blue (stain: 0.1% (w/v) Coomassie blue, 20% (v/v) methanol, 10% (v/v) acetic acid; destain: 50% (v/v) MeOH, 10 % (v/v) acetic acid).

### Bimolecular fluorescence complementation (BiFC)

SPYCE/SPYNE epitope tags for BiFCs (Walter *et al*., 2004) were adapted for Golden Gate cloning. Epitope-tagged interaction candidates were transiently co-expressed in *N. benthamiana*. YFP fluorescence complementation was visually assessed via confocal laser scanning microscopy. For quantification, YFP fluorescence of leaf discs from transformed areas was measured with a plate reader as described (Fischer *et al*., 2017). Protein expression was validated by immunoblot.

### Trypan blue staining

*N. benthamiana* leaf discs (Ø 2 cm) were incubated (5 min, 95°C) in 3 mL trypan blue staining solution (25% (v/v) lactic acid, 25% (v/v) phenol, 25% (v/v) glycerol, 25% (v/v) H_2_O, 25% (w/v) trypan blue). Leaves were washed several times with destaining solution (250 g chloral hydrate in 100 mL H_2_O) and rinsed with water before photo-documentation.

### Elicitors

For elicitation, synthetic 3-OH-C10:0 (Matreya LLC) in MeOH or flg22 (QRLSTGSRINSAKDDAAGLQIA, PEPMIC) in ddH_2_O and corresponding controls were used.

### ROS measurement

ROS was measured with a luminol-based reporter system and a microplate reader (Tecan Infinite F200 PRO or Luminoskan Ascent 2.1, Thermo Fisher Scientific) as described (Ranf *et al*., 2015). Leaf discs (Ø 4 mm) of Brassicaceae (6-8 weeks old) or transiently transformed *N. benthamiana* were floated on water in white 96-well plates for >6 h (*N. benthamiana*) or overnight (Brassicaceae). Prior to measurements, water was replaced with 100 µL HRP/luminol solution (2 μg/mL HRP, 10 μM L-012 (WAKO Chemicals GmbH)). After elicitation, luminescence was recorded in relative light units (RLU) in 1-minute intervals up to 60 minutes. Total ROS accumulation was summed up as indicated.

### Cytosolic calcium measurement

Cytosolic [Ca^2+^] was measured with an aequorin-based reporter system in 9-11 days-old seedlings grown in liquid culture as described (Ranf *et al*., 2012). Seedlings were incubated in 100 µl of 10 μM coelenterazine-h (p.j.k. GmbH) overnight. Luminescence was measured upon elicitation in 10-second intervals for 30 minutes in a microplate luminometer (Luminoskan Ascent 2.1, Thermo Scientific) before discharging. For each time point, luminescence was normalized to total luminescence counts remaining (L/L_max_) and total [Ca^2+^]_cyt_ after elicitation was summed up.

### Purification of LORE-ECDs

ECDs with N-terminal Twin-strep-tag were transiently expressed in *N. benthamiana* apoplasts and harvested via apoplastic washing fluids (AWFs). The ECDs were purified by Strep-tag affinity chromatography (Strep-Tactin®XT 4Flow, IBA Lifesciences, equilibrated in 100 mM Tris pH 8.0, 150 mM NaCl, 1 mM EDTA, with 50 mM biotin for protein elution). Proteins were further purified by size exclusion chromatography (SEC) on a Superdex 200 Increase 10/300 GL (Cytiva), equilibrated in 20 mM Tris pH 8.0, 150 mM NaCl, 1 mM EDTA) and concentrated by filtration (Vivaspin 500 centrifugal concentrator, Sartorius, 30 kDa molecular weight cut-off (MWCO)).

### Ligand depletion binding assay

As described (Shu *et al*., 2021), ECDs were transiently expressed in *N. benthamiana* apoplasts and harvested via AWFs, which were concentrated and incubated with 3-OH-C10:0 (9:1, v/v). Unbound elicitor was separated by filtration (Vivaspin 500 centrifugal concentrator, Sartorius, 30 kDa MWCO). Filtrates were tested for eliciting [Ca^2+^]_cyt_ changes in LORE-OE lines. 5 µL concentrated AWF (1.5 mg/mL total protein concentration) was analyzed for protein expression by immunoblot. Ligand binding assay was performed as described (Shu *et al*., 2021). Purified ECDs (500 nM) were incubated with 3-OH-C10:0 (72 ml:8 ml). Unbound 3-OH-C10:0 was separated by filtration (30 kDa MWCO). Retentates were incubated at 98°C for 30 minutes and filled up to the original volumes. 3-OH-C10:0 released from heat-denatured proteins was separated by filtration (30 kDa MWCO) and used for [Ca^2+^]_cyt_ measurements in LORE-OE lines.

### Red light chlorophyll fluorescence measurement

Red light emission of chlorophyll fluorescence can be measured to quantify cell death (Landeo Villanueva *et al*., 2021). Leaf discs (Ø 4 mm) of transiently transformed *N. benthamiana* were floated on water in black 96-well plates, and chlorophyll fluorescence was measured with a plate reader (Tecan Infinite F200 PRO, excitation 535 nm, emission 590 nm, 25 flashes, integration time 20 µs, 4×4 reads per well, gain 80) as relative fluorescence units (RFU). Values of all reads per well were summed up.

### Microscopy

Confocal laser scanning microscopy was performed using a Leica TCS SP5 (Argon laser, HyD2 detectors) or Olympus FV3000 (diode lasers, PMT detectors). Images were acquired with a size of 512202 cl:192512 pixels. In case of co-expression analysis, images were acquired with a sequential scan to avoid ‘bleed through’. Excitation/emission was detected at 488/500-540 nm for GFP, 561/570-620 nm for mCherry, and 514/525-550 nm for YFP. Images were processed with OMERO (Allan *et al*., 2012).

### Fluorescence lifetime imaging (FLIM)

GFP fluorescence lifetimes were measured via time-correlated single-photon counting (TCSPC) by an Olympus FV3000 system linked to a PicoQuant FCS/FLIM-FRET/rapidFLIM upgrade kit, based on described protocols (Weidtkamp-Peters & Stahl, 2017). TCSPC was performed with a 485 nm (LDH-D-C-485) pulsed laser, two TCSPC modules (TimeHarp 260 PICO Dual, TimeHarp 260 NANO Dual) and two-photon counting PMA hybrid 40 detectors. Co-expressing cells were detected by confocal imaging via an Olympus FV3000 60x water immersion objective (UPLSAPO60XW 60x/NA 1.2/WD 0.28), and a selected measuring area was magnified (4x zoom). TCSPC was performed with a laser pulse rate of 40.00 MHz, TCSPC resolution of 25.0 ps and image size of 512206 cl:772512 pixels. 500-1000 photon counts per pixel were acquired per image. FLIM was analyzed using Symphotime64 software (PicoQuant) via n-exponential reconvolution and an internally calculated instrument response function (IRF). A two-parameter fitting (n=2) was performed in most cases. Intensity-weighted average lifetimes τ of regions of interest containing membranes were determined for each FLIM image. FLIM analyses with fitting coefficients (χ^2^) between 1.0 and 2.0 were accepted.

## RESULTS

### *At*LORE forms receptor homomers *in planta*

*At*LORE and putative LORE orthologues from Brassicaceae (**Fig S1A**) are structurally related to well-studied, homomerizing *Bra*SRKs. High protein sequence similarity and conservation of disulfide bridge-forming cysteine residues indicate a similar ECD architecture of *At*LORE and *Bra*SRKs (**Fig S1B, C**), which is supported by a structural alignment of the *Bra*SRK9-ECD crystal structure and the predicted *At*LORE-ECD model (**Fig S1D**). This suggests that *At*LORE might form receptor homomers like BraSRKs. Homomerizing RLKs comprising a strong RD-type kinase domain can be prone to spontaneous auto-activation, which is often associated with overshooting signaling and cell collapse. Indeed, transient overexpression of *At*LORE in *Nicotiana benthamiana* results in a cell death-like necrotic phenotype (**Fig S2A**), which can be visualized by trypan blue staining (**Fig S2B**) and increased red light emission upon chlorophyll excitation (**Fig 1A**) (Landeo Villanueva *et al*., 2021). Mutation K516A in the conserved ATP binding site (kinase-mutated, Km; **Fig S1E**) renders the *At*LORE kinase domain inactive (Ranf *et al*., 2015; Luo *et al*., 2020) and abolishes cell death (**Fig 1A, S2**), indicating that cell death depends on active kinase signaling. This may suggest that *At*LORE is auto-activated upon overexpression in *N. benthamiana* by spontaneous, ligand-independent receptor homomerization, further supporting our initial hypothesis. Thus, we analyzed protein-protein interactions *in vivo*. Co-immunoprecipitation (Co-IP) of kinase-active and kinase-mutated *At*LORE supports homomerization upon transient expression in *N. benthamiana* (**Fig 1B**). *At*LORE is less abundant than *At*LORE-Km, likely due to ongoing cell death and associated protein degradation. To avoid protein instability and increased background auto-fluorescence caused by cell death, *At*LORE-Km was used for all fluorescence-based interaction assays. Self-interaction of *At*LORE-Km was verified by bimolecular fluorescence complementation (BiFC) (**Fig S3A-C**) and Förster resonance energy transfer (FRET) fluorescence lifetime imaging (FLIM) in *N. benthamiana* (**Fig 1C and Fig S3D**). Co-expression of *At*LORE-Km-mCherry with *At*LORE-Km-GFP reduces GFP fluorescence lifetime τ, while co-expression of mCherry-tagged GLUTATHIONE-S TRANSFERASE (*At*GST-mCherry) does not. FLIM images show equally distributed protein interaction along the membrane (**Fig S3D**). *At*LORE homomerization does not significantly change upon treatment with the ligand 3-OH-C10:0 (**Fig 1D**). Thus, under the experimental conditions used, *At*LORE forms homomers in the plasma membrane and auto-activates in a ligand-independent manner.

**Figure 1.**
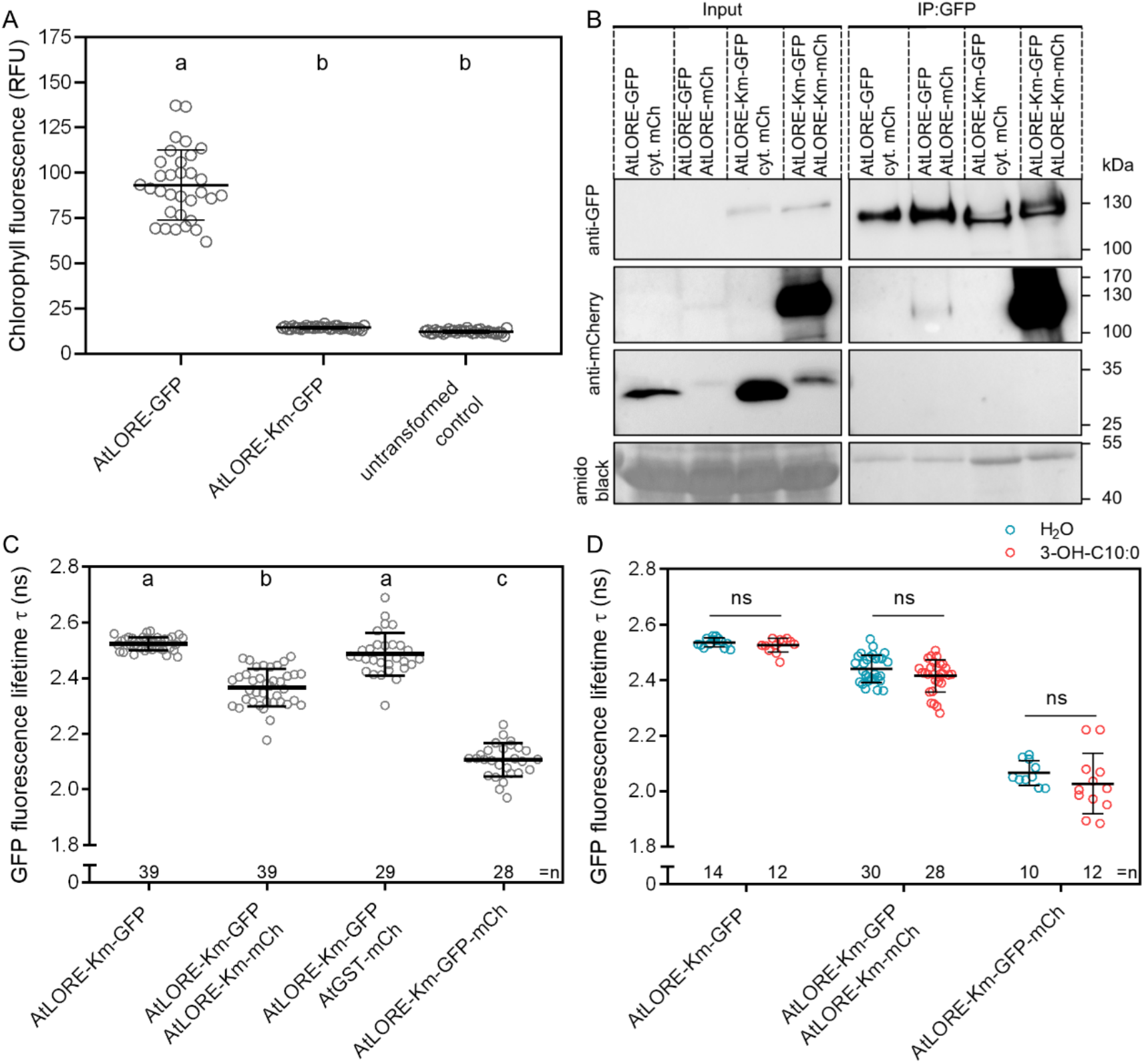
*At*LORE forms homomers and has an auto-immunity phenotype in *Nicotiana benthamiana.* **A** Chlorophyll fluorescence (in relative fluorescence units, RFU) upon transient expression (CaMV35S promoter) of candidates in *N. benthamiana* four days post agro-infiltration. Mean and SD of pooled data of two biological replicates are shown (n=32 leaf discs). Data not sharing the same letter are significantly different as analyzed by one-way ANOVA with Tukey’s multiple comparisons test, α=0.01. **B** Anti-GFP and anti-mCherry (mCh) immunoblot of co-immunoprecipitation (GFP-trap) after transient co-expression (CaMV35S promoter) of interaction candidates in *N. benthamiana* two days post agro-infiltration. Total protein was stained with amido black. Km, kinase-mutated; cyt., cytosolic. **C, D** FRET-FLIM of interaction candidates transiently co-expressed (estradiol-inducible XVE promotor) in *N. benthamiana*. Pooled data of five (C) or two (D) independent biological replicates are shown. Data show means with SD of GFP fluorescence lifetimes τ in nanoseconds (ns); n, number of analyzed cells; mCh, mCherry; ns, not significant. Statistics was analyzed by one-way ANOVA with Tukey’s multiple comparisons test, α=0.01 (C) or two-way ANOVA with Sidak’s multiple comparisons test comparing treatments; P>0.1234 (D). FLIM images (D) were acquired 10-20 minutes after treatment with 5 µM 3-OH-C10:0 or ddH_2_O as control.

### *At*LORE homomerization is mediated by the extracellular and transmembrane domains

To identify the region of *At*LORE mediating its homomerization, we generated truncated variants of *At*LORE containing different combinations of ECD, TMD or ICD (**Fig 2A, S1F, Supplementary Information Table S3**). In Co-IP assays upon co-expression in *N. benthamiana*, full-length *At*LORE-Km-HA strongly associates with full-length *At*LORE-Km-GFP and *At*LORE-ECD-TMD-GFP, but notably less with *At*LORE-TMD-ICD-Km-GFP. In contrast, interaction with apoplastic *At*LORE-ECD-GFP or cytosolic *At*LORE-ICD-Km-GFP was barely detectable, and no interaction was observed with cytosolic GFP (**Fig 2B**). This indicates that primarily ECD and TMD contribute to *At*LORE homomerization. In FRET-FLIM, both *At*LORE-ECD-TMD-mCherry and *At*LORE-TMD-ICD-Km-mCherry reduce the GFP fluorescence lifetime of the FRET-donor *At*LORE-Km-GFP (**Fig 2C and S4**). Furthermore, *At*LORE-ECD-TMD interacts with *At*LORE-TMD-ICD (**Fig 2D**), where only the TMDs can mediate the interaction. While *At*LORE-TMD-ICD-Km-mCherry was partially miss-localized (**Fig S4**), the reduction in GFP fluorescence lifetime of the FRET donor indicates co-localization of the interaction candidates in the plasma membrane. In summary, our data support the contribution of both ECD and TMD to *At*LORE homomerization.

**Figure 2.**
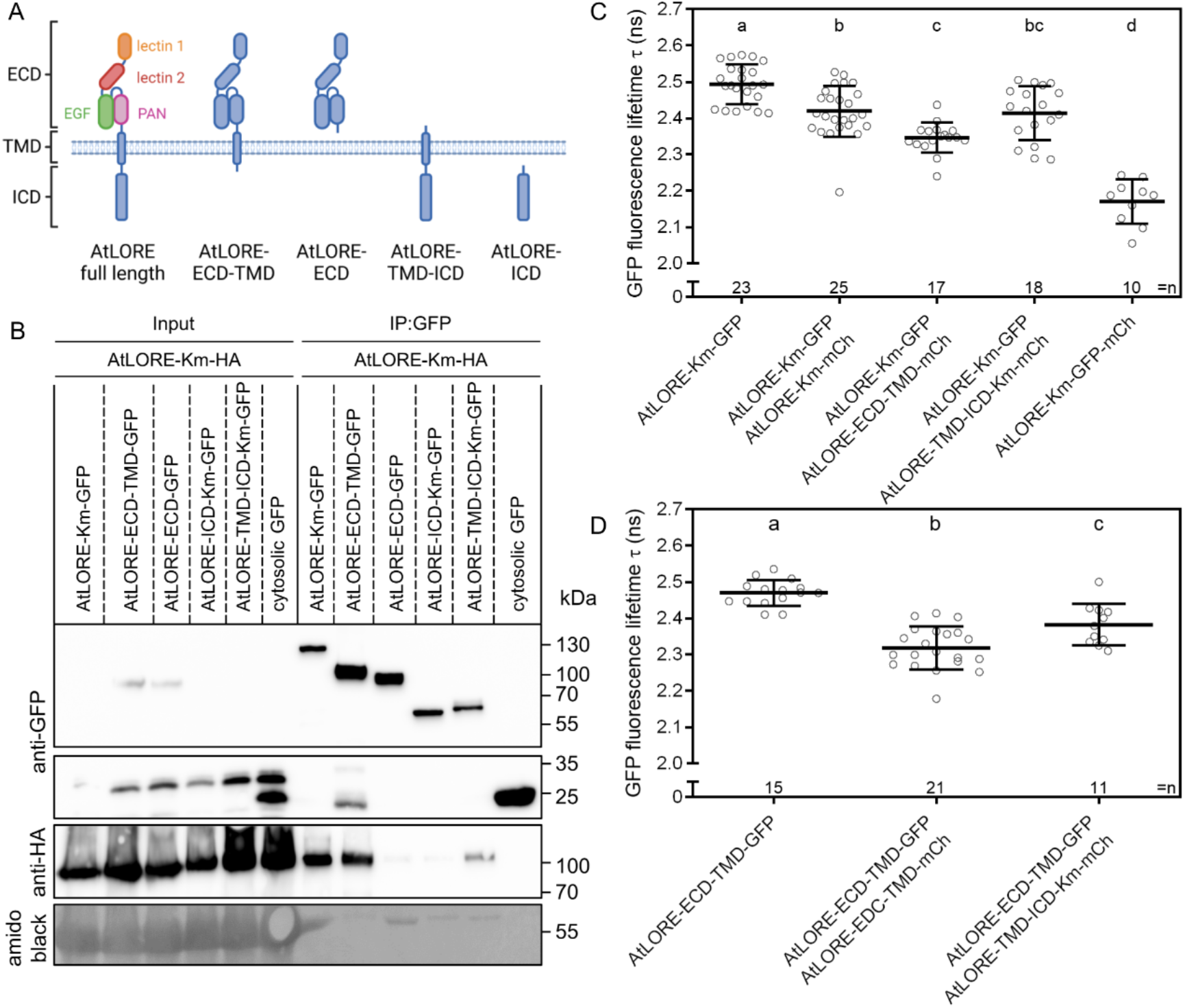
Homomerization of *At*LORE is mediated by the extracellular and transmembrane domains in *Nicotiana benthamiana*. **A** Scheme of *At*LORE truncation variants; ECD, extracellular domain; TMD, transmembrane domain; ICD, intracellular domain. For detailed sequence information, see Supplementary Information Table S3 and Fig S1F. **B** Anti-GFP and anti-HA immunoblot of co-immunoprecipitation (GFP-trap). Full-length and truncated *At*LORE-Km interaction candidates transiently co-expressed (CaMV35S promoter) in *N. benthamiana* were analyzed two days post agro-infiltration. Total protein is stained with amido black. **C, D** FRET-FLIM of full-length *At*LORE-Km-GFP or *At*LORE-ECD-TMD-GFP versus truncated *At*LORE-mCherry variants transiently expressed (estradiol-inducible XVE promotor) in *N. benthamiana*. Pooled data from two independent biological replicates are shown. Data show mean with SD of GFP fluorescence lifetimes τ. n, number of analyzed cells; mCh, mCherry. Data not sharing the same letter are significantly different as analyzed by one-way ANOVA with Tukey’s multiple comparisons test, α=0.01.

### *A. lyrata* and *A. halleri* possess LORE orthologues but do not respond to 3-OH-C10:0

Phylogenetic analysis shows that LORE is restricted to Brassicaceae (Ranf et al., 2015) (**Fig S1A**). Putative LORE orthologues were identified in *Arabidopsis halleri* (Araha.6790s0007.1, *Ahal*LORE), *Arabidopsis lyrata* (AL2G14950, *Alyr*LORE) and *Capsella rubella* (CARUB_v10021901mg, *Crub*LORE) based on protein sequence similarity (Supplementary Information **Fig S1A-B; Table S5**). While the application of 3-OH-C10:0 application triggers the production of reactive oxygen species (ROS), a typical PTI response (Kutschera *et al*., 2019), in leave discs of *A. thaliana* and *C. rubella*, this is not the case for leaf discs of *A. lyrata* and *A. halleri* (**Fig 3A**), while all four species respond to flg22 treatment (**Fig S5A**). For further analysis, RNA from untreated leaf material of *C. rubella*, *A. lyrata* and *A. halleri* was transcribed into cDNA to clone the *LORE* coding sequences (CDS). Full-length CDS of all orthologues were obtained, confirming *LORE* expression in the *A. lyrata* and *A. halleri* leaf tissue used for ROS assays (**Fig 3A, S5B**). The amino acid sequence of *Ahal*LORE slightly varied from the publicly available database sequence (**Fig S6**). All GFP-tagged LORE orthologues are expressed and localize to the plasma membrane in *N. benthamiana* (**Fig S5B**). Solanaceous *N. benthamiana* can gain the function of 3-OH-C10:0 sensing by transient expression of *At*LORE early after agro-infiltration and before the onset of macroscopic cell death (Ranf *et al*., 2015; Kutschera *et al*., 2019). Cloned LORE orthologues were tested in gain-of-function (GOF) ROS assays in *N. benthamiana* (**Fig 3B**). Expression of *At*LORE or *Crub*LORE, but not *Ahal*LORE or *Alyr*LORE, results in ROS production upon 3-OH-C10:0 treatment. We generated stable complementation lines expressing *At*LORE, *Alyr*LORE and *Crub*LORE in the *lore*-1 knock-out background under the endogenous promotor (p*At*LORE) (Ranf *et al*., 2015) to assess their functionality in *A. thaliana*. Both, ROS measurements on soil-grown plants and [Ca^2+^]_cyt_ measurements in seedlings show full complementation of 3-OH-C10:0 signaling for *At*LORE and *Crub*LORE, but no or only partial complementation for *Alyr*LORE (**Fig 3C-D**). Taken together, the inability of *A. lyrata* and *A. halleri* to respond to 3-OH-C10:0 can be attributed to their poorly functioning LORE orthologues.

**Figure 3.**
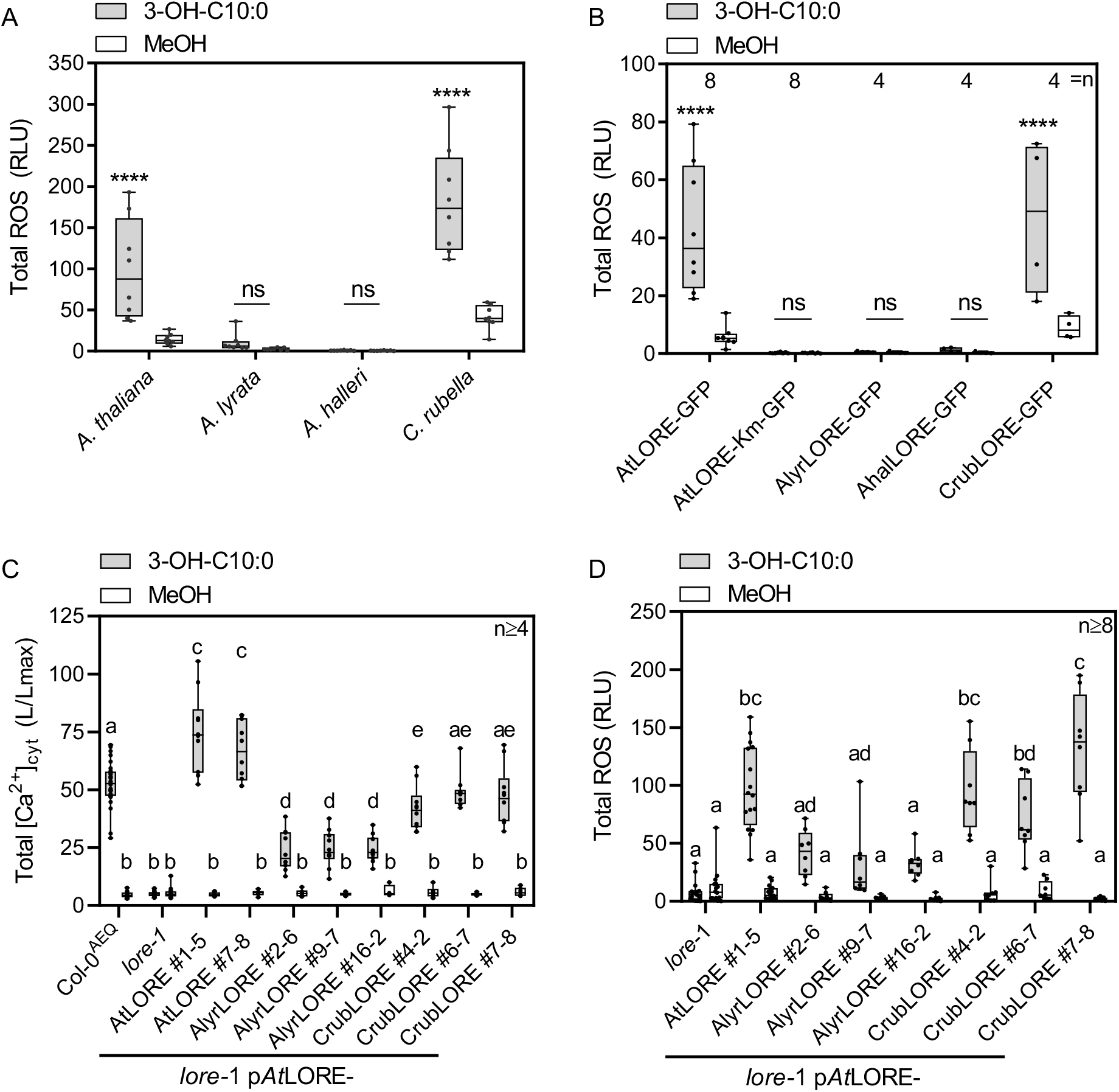
3-OH-C10:0-induced immune-responses of *Arabidopsis lyrata*, *Arabidopsis halleri* and *Capsella rubella* LORE orthologues. **A** Total ROS accumulation of *A. halleri*, *C. rubella*, *A. lyrata* and *A. thaliana* leaf discs treated with 5 µM 3-OH-C10:0 or MeOH as control. Median with minimum to maximum of total ROS between 3-60 minutes after elicitation from one biological replicate is shown; n=8 leaf discs. Measurements were repeated at least two times with similar outcomes. **B** Total ROS accumulation of *N. benthamiana* leaf discs transiently overexpressing (CaMV35S promoter) *At*LORE or putative LORE orthologues upon application of 1 µM 3-OH-C10:0 or MeOH as control. Median with minimum to maximum of total ROS between 3-45 minutes after treatment is shown; n, number of leaf discs. Data of two technical replicates are shown. **A, B** Significant difference between MeOH and 3-OH-C10:0 treatment was tested by two-way ANOVA with Sidak‘s multiple comparisons test; ****, P≤0.0001; ns, not significant, P>0.1234. **C, D** [Ca^2+^]_cyt_ measurements in seedlings (C) and ROS accumulation of soil-grown plants (D) from stable transgenic complementation lines (*At*LORE promoter) of different LORE orthologous in the *lore-*1 background. Samples were elicited with 1 µM 3-OH-C10:0 or a MeOH control. Data from up to three technical replicates performed in two independent biological replicates were pooled. ROS data were normalized to the average background signal. Significant difference was tested by two-way ANOVA with Tukey’s multiple comparisons test; data not sharing the same letters are significantly different, α=0.01

### *Alyr*LORE has signaling and elicitor binding capacity

To identify the dysfunctional domain of *Alyr*LORE, we created reciprocal domain swaps (DS) between *At*LORE and *Alyr*LORE and tested them for GOF-ROS responses in *N. benthamiana* (**Fig 4A-B, S1F**, **Supplementary Information Table S3**). Replacing the ECD and TMD of *At*LORE with *Alyr*LORE renders the chimera inactive, while swapping the *At*LORE-ICD with *Alyr*LORE-ICD keeps it functional. Thus, *Alyr*LORE has a signaling-competent ICD, but a defective ECD. To test whether *Alyr*LORE-ECD is impaired in ligand binding, we performed ligand-depletion experiments, which we had previously used to show that *At*LORE and *Crub*LORE can bind 3-OH-C10:0, while the closely related *At*SD1-23 cannot (Ranf *et al*., 2015; Shu *et al*., 2021). ECDs expressed in the apoplast of *N. benthamiana* leaves were harvested in apoplastic washing fluids. Equal amounts of total protein (**Fig S7A**) were incubated with 3-OH-C10:0. Filtration of receptor-ligand mixtures through membranes with a 30 kDa cut-off results in the retention of receptor and bound ligand. Unbound 3-OH-C10:0 in the filtrates is detected by [Ca^2+^]_cyt_ measurements in *A. thaliana* seedlings overexpressing (OE) *At*LORE. The ECDs of all tested LORE orthologues bind and completely deplete 3-OH-C10:0 from filtrates, as no [Ca^2+^]_cyt_ response is detectable (**Fig 4C**). To compare ligand-binding capacities of *At*LORE and *Alyr*LORE, we performed in-vitro ligand-binding assays in a concentration-dependent manner with ECDs purified via Streptavidin-tag affinity and size exclusion chromatography (**Fig S7B-D**). The procedure of the depletion assay was repeated with purified ECDs and varying concentrations of 3-OH-C10:0, but this time retentates were heat-denatured to release bound 3-OH-C10:0 which was subsequently used to elicit *At*LORE-OE seedlings. *At*LORE and *Alyr*LORE ECDs show similar 3-OH-C10:0 binding capacities at all tested concentrations (**Fig 4D**). Since neither kinase activity nor ligand binding of *Alyr*LORE is affected, the impaired 3-OH-C10:0 perception of *Alyr*LORE must have other causes.

**Figure 4.**
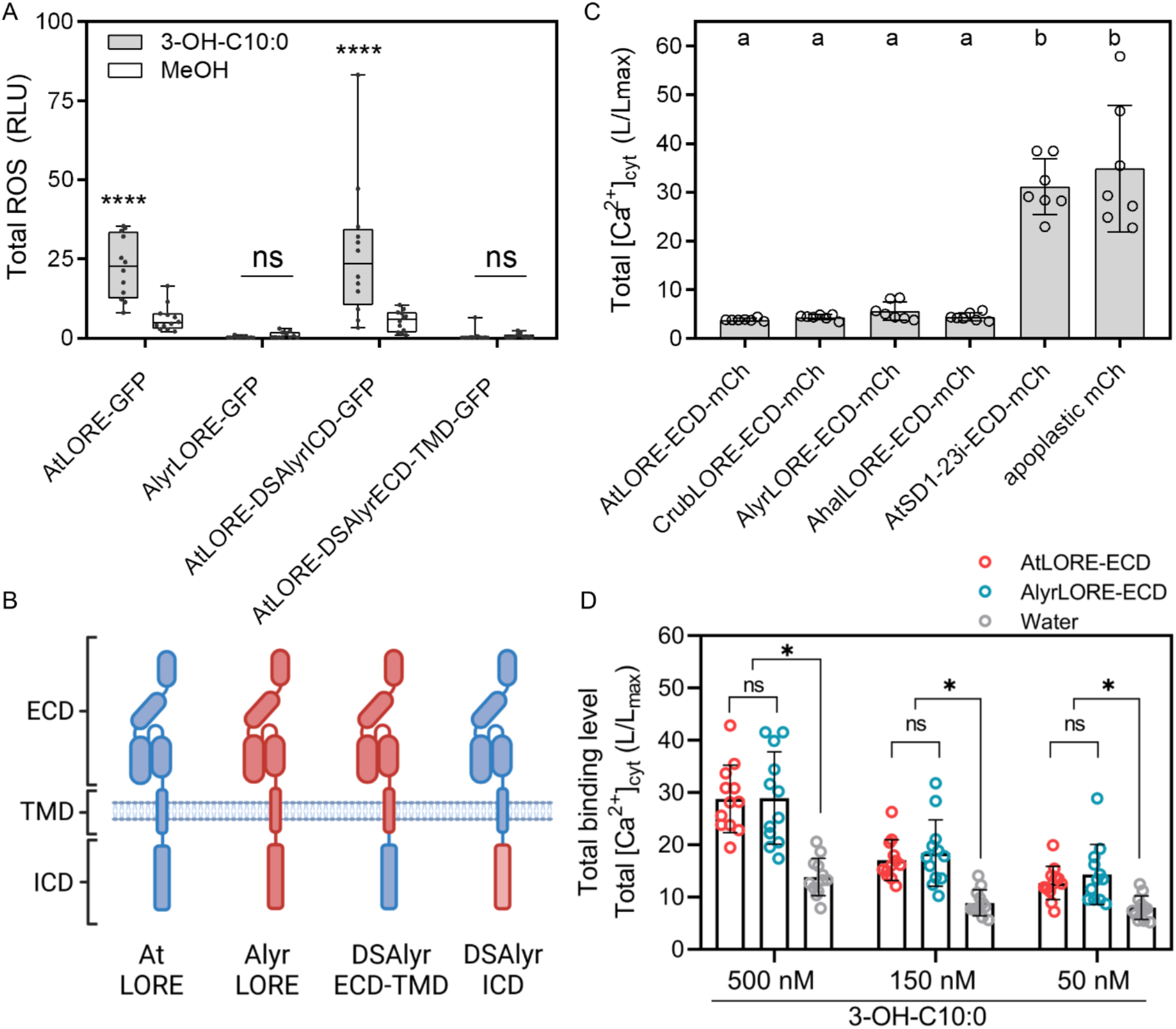
A defect in the *Alyr*LORE ectodomain is responsible for the lack of 3-OH-C10:0 perception in *Nicotiana benthamiana*, but the ability to bind the ligand is not impaired. **A** Total ROS accumulation in *N. benthamiana* transiently expressing (CaMV35S promoter) *At*LORE, *Alyr*LORE and chimera with either ECD-TMD or ICD domain swap (DS) from 1-41 minutes after elicitation with 10 µM 3-OH-C10:0 or MeOH. Median with minimum to maximum is shown; n=12 leaf discs. Statistics were analyzed by two-way ANOVA with Sidak’s multiple comparisons test; ****, P≤0.0001; ns, not significant, P>0.1234. **B** Scheme of *At*LORE-*Alyr*LORE domain swaps; ECD, extracellular domain; TMD, transmembrane domain; ICD, intracellular domain. For detailed sequence information and domain swap sites, see Supplementary Information Table S3 and Fig S1F. **C** 3-OH-C10:0 binding to LORE-ECDs was tested in a ligand-depletion assay. Unbound 3-OH-C10:0 in filtrates was detected by [Ca^2+^]_cyt_ measurements of LORE-overexpression (OE) lines. Full depletion of the [Ca^2+^]_cyt_ response indicates ligand binding. The ECD of the *At*LORE paralog *At*SD1-23 and apoplastic mCherry were used as controls; i, integrated LORE intron. Pooled data from two independent biological replicates are depicted. Data show mean and SD of total [Ca^2+^]_cyt_ from 0.5 to 30 minutes after elicitation; n=7 seedlings. Data not sharing the same letter are significantly different as analyzed by one-way ANOVA with Tukey’s multiple comparisons test, α=0.01. **D** Ligand binding assay was performed with 500 nM purified ECDs of *At*LORE, *Alyr*LORE, or water and different 3-OH-C10:0 concentrations. Bound 3-OH-C10:0 released from heat-denatured ECDs was detected by [Ca^2+^]_cyt_ measurements of LORE-OE lines. Pooled data from four independent biological replicates are depicted. Data show mean and SD of total [Ca^2+^]_cyt_ from 0.5 to 30 minutes after elicitation; n=12 seedlings. Significant difference was tested by two-way ANOVA with Tukey’s multiple comparisons test; α=0.1. *, P≤0.1; ns, not significant, P>0.1.

### LORE orthologues from 3-OH-C10:0 non-responsive species cannot homomerize

Unlike *At*LORE or *Crub*LORE, overexpression of *Alyr*LORE or *Ahal*LORE in *N. benthamiana* does not cause cell death, as determined by chlorophyll fluorescence measurements (**Fig 5A**). We hypothesized that *Alyr*LORE and *Ahal*LORE fail to auto-activate due to impaired homomerization. Indeed, *Alyr*LORE and *Ahal*LORE are unable to homomerize in FRET-FLIM in *N. benthamiana,* in contrast to *Crub*LORE (**Fig 5B-D).** *Alyr*LORE also does not interact with *At*LORE (**Fig 5B**). LORE homomerization thus appears to be crucial for the activation of signaling. We conclude that the cell death phenotype in *N. benthamiana* upon LORE overexpression serves as an indicator of LORE signaling functionality and requires both kinase activity and receptor homomerization. As *Alyr*LORE and *Ahal*LORE bind 3-OH-C10:0 but do not homomerize, ligand binding occurs independently of receptor homomerization. In support of this, both *At*LORE- and *Alyr*LORE-ECDs elute as single peaks during SEC, which suggests a monomeric state of the soluble ECDs, and both purified ECDs bind 3-OH-C10:0 (**Fig 4D, Fig S7**).

**Figure 5.**
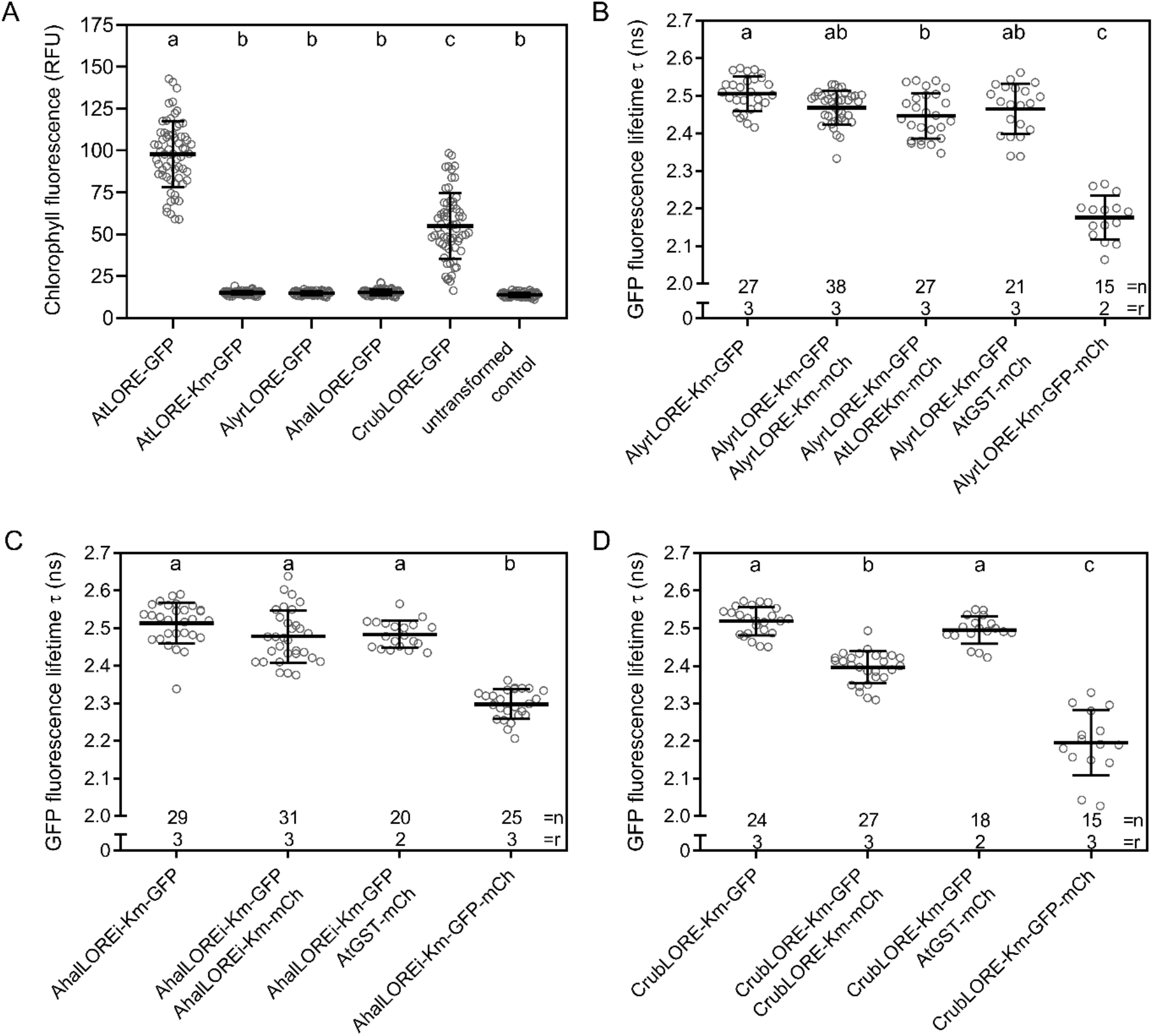
LORE orthologues from 3-OH-C10:0 non-responsive species are impaired in homomerization in *Nicotiana benthamiana*. **A** Chlorophyll fluorescence measurement (in relative fluorescence units, RFU) of *N. benthamiana* leaf discs transiently overexpressing (CaMV35S promoter) different LORE orthologues four days post agro-infiltration. Mean and SD of pooled data from two biological replicates with two or three technical replicates are shown; n=64 leaf discs. Statistics were analyzed by one-way ANOVA with Tukey‘s multiple comparisons test, α=0.01. Data not sharing the same letter are significantly different. **B-D** FRET-FLIM of putative LORE orthologues from *A. lyrata* (**B**), *A. halleri* (**C**) and *C. rubella* (**D**) transiently expressed (estradiol-inducible XVE promotor) in *N. benthamiana*. Km, kinase-mutated; i, integrated *At*LORE intron; n, number of analyzed cells; r, number of biological replicates; mCh, mCherry. Data show mean with SD of GFP fluorescence lifetimes τ of pooled data of biological replicates as indicated. Statistics were analyzed by one-way ANOVA with Tukey‘s multiple comparisons test, α=0.01. Data not sharing the same letters are significantly different.

### Mapping of homomerization region using *At*LORE and *Alyr*LORE chimera

While *Alyr*LORE and *Ahal*LORE share a very high overall amino acid identity with *At*LORE, they have several single amino acid polymorphisms (SAPs) in their ECDs compared to the *At*LORE-ECD, distributed across all domains (**Fig S1F**). To identify the region of the ECD causing the loss of homomerization, we generated ECD and TMD chimera between *At*LORE and *Alyr*LORE or exchanged individual domains, such as lectin 1, lectin 2, EGF or PAN domain (L1, L2, E, P; **Fig S8A**) and combinations of domains (LL, EP; **Fig 6A**). After verifying the expression and membrane localization of the chimera in *N. benthamiana* (**Fig S9**), we analyzed homomerization via FRET-FLIM (**Fig 6 and S8B**). Since all chimera with individual domain swaps retained the ability to homomerize (**Fig S8B**), the loss of *Alyr*LORE homomerization is not caused by a single domain. Therefore, we tested different domain combinations (**Fig 6B, S8C**). Interestingly, only substituting the whole ECD of *At*LORE by *Alyr*LORE, but not partial ECD (LL, EP) or TMD-ICD swaps, impairs homomerization in FRET-FLIM (**Fig 6B, S8C**). To rule out that cross-connections between LL and EP domains support homomerization, we tested LL and EP domain swaps against each other, but homomerization remained functional (**S8C**). Similar results were observed in 3-OH-C10:0-triggered ROS accumulation or chlorophyll fluorescence assays upon overexpression of these chimera in *N. benthamiana* (**Fig 4A, 6C and D**). Overall, chimera with partial ECD swaps remain signaling-competent, while those with ECD-TMD exchange do not. We could not pin down a specific region of the *Alyr*LORE-ECD or TMD that causes the loss of homomerization, which may indicate a large, domain-spanning interaction interface of *At*LORE.

**Figure 6.**
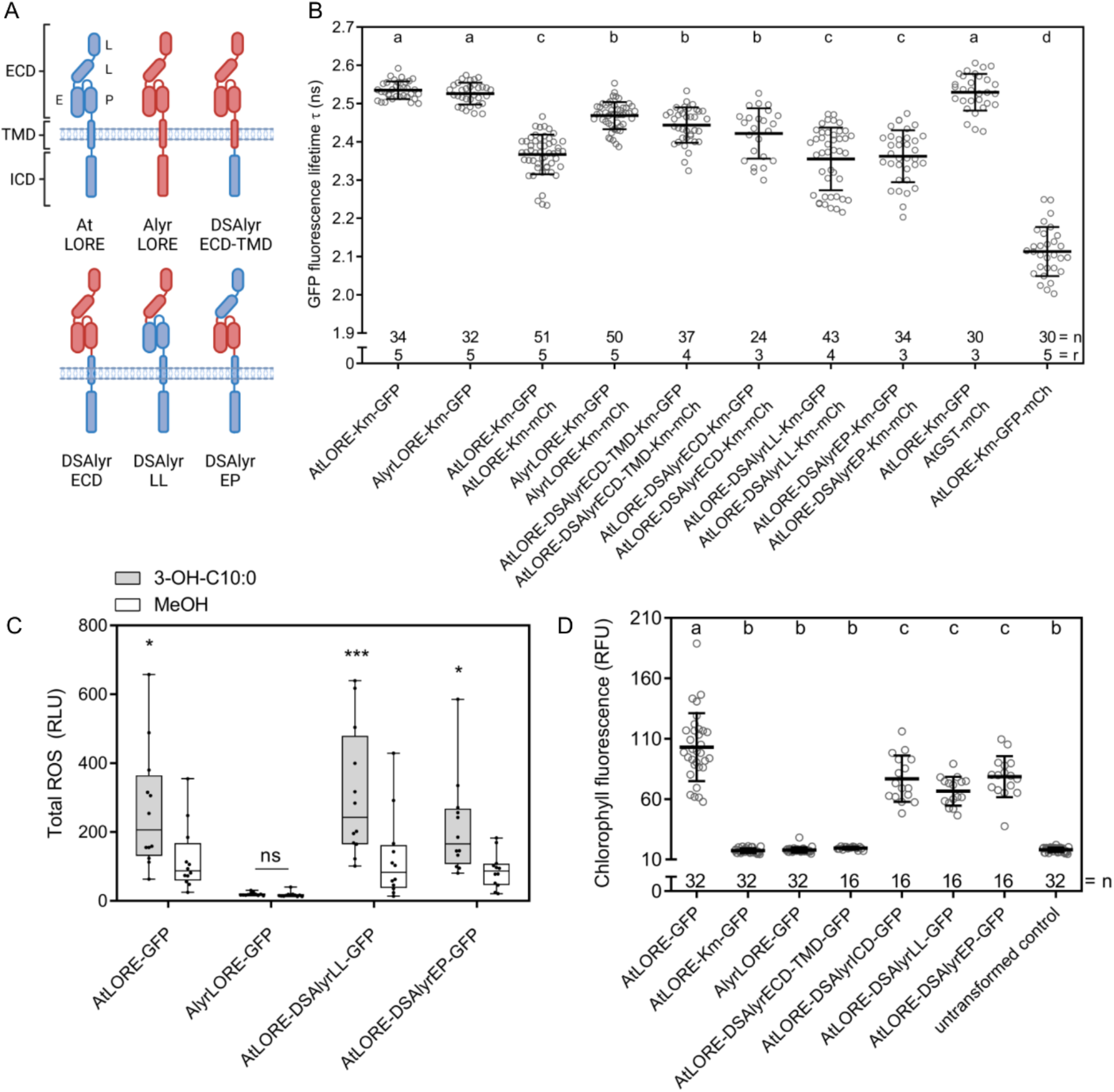
Mapping of the dimerization region using receptor chimera of *At*LORE and *Alyr*LORE in *Nicotiana benthamiana*. **A** Schematic overview of chimera between *Alyr*LORE and *At*LORE. DS, domain swap; L, lectin domain; E, EGF domain; P, PAN domain. **B** FRET-FLIM of *At*LORE-*Alyr*LORE domain swaps (DSAlyr) transiently expressed (estradiol-inducible XVE promotor) in *N. benthamiana*. Mean and SD of GFP fluorescence lifetimes τ of pooled data are shown; r, number of biological replicates; n, number of analyzed cells. Significant difference between MeOH and 3-OH-C10:0 treatment was tested by two-way ANOVA with Sidak‘s multiple comparisons test, α=0.01. **C** Gain-of-function ROS measurements of *N. benthamiana* transiently expressing (CaMV35S promoter) *At*LORE-*Alyr*LORE chimera 1-44 minutes after elicitation with 5 µM 3-OH-C10:0 or MeOH as control. Graph shows median with minimum to maximum; n=12 leaf discs. Statistics were analyzed by two-way ANOVA with Sidak’s multiple comparisons test, *, P≤0.0332; ***, P≤0.0002. Experiment was repeated two times with similar outcome. **D** Chlorophyll fluorescence measurements (in relative fluorescence units, RFU) in *N. benthamiana* upon transient expression (CaMV35S promoter) of *At*LORE-*Alyr*LORE chimera four days post agro-infiltration. Graph shows mean with SD of one biological replicate. Experiment was repeated twice with similar outcome; n, number of leaf discs. Statistics were analyzed by one-way ANOVA with Tukey‘s multiple comparisons test, α=0.01. Data not sharing the same letter are significantly different.

### Homomerization is required for *At*LORE downstream signaling

The LORE kinase domain auto-phosphorylates *in vitro* (Luo *et al*., 2020). Our data suggest that LORE homomerization is required for its activation and downstream signaling. Therefore, we performed competition experiments with the truncated *At*LORE-ECD-TMD variant (**Fig 7**), which can form hetero-complexes with full-length *At*LORE (**Fig 2B and C**). These dimers are presumably signaling-incompetent since one monomer lacks a kinase domain. To outcompete full-length *At*LORE homo-complexes, *At*LORE-ECD-TMD must be present in excess relative to *At*LORE. We exploit the intrinsically high protein accumulation of *At*LORE-ECD-TMD compared to full-length *At*LORE upon transient expression in *N. benthamiana* (**Fig 2B, S10**) and increased the ratio of Agrobacteria carrying the competitor expression plasmid to enhance the effect. To prevent competition for ligand binding, we used an excess of ligand (5 µM 3-OH-C10:0). Indeed, when co-expressed in excess in *N. benthamiana*, *At*LORE-ECD-TMD outcompetes signaling-competent full-length *At*LORE homomers, as shown by a fully suppressed ROS response upon 3-OH-C10:0 elicitation, while cytosolic mCherry does not (**Fig 7A**). *At*LORE-TMD-ICD, *Alyr*LORE-ECD-TMD or *At*LORE-ECD are not able to suppress 3-OH-C10:0 induced *At*LORE ROS signaling (**Fig 7B and C**). Similarly, *At*LORE-ECD-TMD reduces the cell death caused by *At*LORE overexpression in *N. benthamiana*, while *Alyr*LORE-ECD-TMD or *At*LORE-ECD do not (**Fig 7D**). All truncated proteins accumulated to a similar extent (**Fig S10**). To assess whether we can suppress 3-OH-C10:0-induced signaling of endogenous *At*LORE in Arabidopsis, we generated stable transgenic Col-0 aequorin (Col-0^AEQ^) lines that overexpress either *At*LORE-ECD-TMD-mCherry or *At*LORE-ECD-mCherry. Expression and localization of the stably overexpressed *At*LORE truncations in these lines were confirmed by microscopy (**Fig S11**). In [Ca^2+^] measurements, 3-OH-C10:0-induced calcium signaling is strongly decreased in lines expressing the homomerization-outcompeting variant *At*LORE-ECD-TMD but remains on wild-type level for lines expressing soluble *At*LORE-ECD in the apoplast (**Fig 7E**). In conclusion, the observed dominant negative effect of overexpressed *At*LORE-ECD-TMD on *At*LORE signaling in both *N. benthamiana* and *A. thaliana* is caused by the formation of signaling-incompetent receptor complexes and supports that *At*LORE homomerization is essential for 3-OH-C10:0 induced immune signaling.

**Figure 7.**
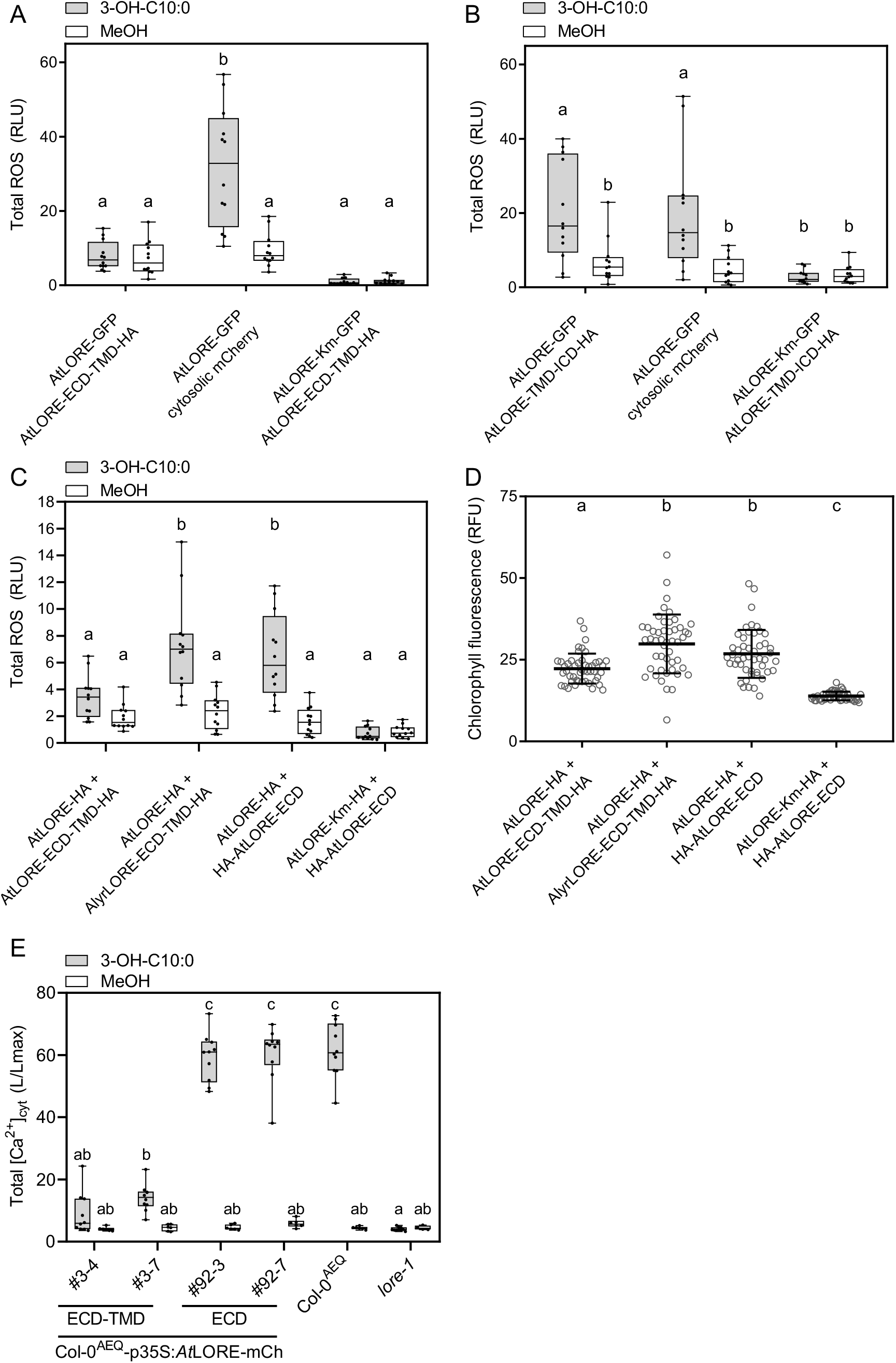
*At*LORE homomerization is required for downstream signaling in *Nicotiana benthamiana* and *Arabidopsis thaliana*. **A, B** Dominant negative effect of co-expressed (CaMV35S promoter) *At*LORE truncations or cytosolic mCherry on total ROS accumulation of full-length *At*LORE in *N. benthamiana* upon elicitation with 5 µM 3-OH-C10:0 or MeOH as a control. Median with minimum to maximum of total ROS from 3-45 minutes is shown; n=12 leaf discs. Three biological replicates were performed with similar outcomes. **C, D** Dominant negative effect of co-expressed (CaMV35S promoter) *At*LORE or *Alyr*LORE truncations on total ROS accumulation (C) or chlorophyll fluorescence (D) in *N. benthamiana*. Both readouts were acquired from the same set of three independent agro-infiltration samples; pooled data of all biological replicates are shown. Gain-of-function ROS accumulation (C) was measured 30-36 hours post agro-infiltration upon elicitation with 5 µM 3-OH-C10:0 or MeOH. Median with minimum to maximum of total ROS accumulation from 4-59 minutes after elicitation is shown; n=12 leaf discs. Chlorophyll fluorescence (D) was measured four days after agro-infiltration. Mean with SD of pooled data are shown; n=48 leaf discs. **E** [Ca^2+^]_cyt_ was measured on seedlings of Col-0^AEQ^ reporter lines stably overexpressing (CaMV35S promoter) either *At*LORE-ECD-TMD-mCherry or *At*LORE-ECD-mCherry. FastRed-selected seedlings of the segregating T2 generation of two independent lines were elicited with 5 µM 3-OH-C10:0 or MeOH. Pooled data from two independent biological replicates are depicted. Data show mean and SD of total [Ca^2+^]_cyt_ from 0 to 30 minutes after elicitation; n=10 seedlings for 3-OH-C10:0; n≥4 for MeOH; mCh, mCherry. Statistics were analyzed by two-way (A-C,E) or one-way (D) ANOVA and Tukey’s multiple comparisons test, α=0.01. Data not sharing the same letter are significantly different.

## DISCUSSION

As the first SD-RLK from *A. thaliana* with a known ligand and characterized function (Ranf *et al*., 2015; Kutschera *et al*., 2019), *At*LORE is an important model for studying signaling mechanisms of SD-RLKs. Yet, the mechanism of *At*LORE receptor activation remained largely unknown. Numerous examples of receptor hetero-dimerization have been found in plants, making it the predominant concept of receptor activation mechanisms (Burkart & Stahl, 2017; Wan *et al*., 2019; Gou & Li, 2020). In contrast, the receptor activation by homomerization, as observed here for *At*LORE (**Fig 1**), seems rarer but common for representatives of the SD-RLK family. The SD-RLKs LARGE SPIKE S-DOMAIN RECEPTOR LIKE KINASE 1 (*Os*LSK1) from rice homomerizes and heteromerizes with five close homologs *in yeast* and *in planta* (Zou *et al*., 2015). Homo-dimerization of *Bra*SRKs has been studied in more detail and is required for ligand binding and signal transduction in SI (Giranton *et al*., 2000; Naithani *et al*., 2007; Shimosato *et al*., 2007; Ma, R *et al*., 2016; Murase *et al*., 2020). *Bra*SRKs seem to pre-assemble into homo-dimers, which rearrange upon ligand binding to enhance *Bra*SRK dimerization, thus allowing rapid receptor activation (Giranton *et al*., 2000; Naithani *et al*., 2007; Shimosato *et al*., 2007). In some species, S-locus glycoproteins (SLGs), which are secreted and highly homologous to their corresponding SRK-ECD, enhance SI by forming a receptor complex with SRKs, but without having direct ligand binding affinity (Takasaki *et al*., 2000; Takayama *et al*., 2001; Shimosato *et al*., 2007; Nasrallah, 2023). The detailed molecular mechanism of how SLGs are involved in SI remains unclear.

Whether preformed receptor complexes exist prior to ligand binding under physiological conditions is controversially discussed for several RLKs. *At*CERK1 forms homo-dimers independent of chitin binding upon overexpression in *A. thaliana* (Liu *et al*., 2012), whereas *At*CERK1 homo-dimerization was fully dependent on *At*LYK5 and chitin binding under physiological conditions (Cao *et al*., 2014; Gong *et al*., 2020). In *A. thaliana*, the LRR-RLK BRASSINOSTEROID INSENSITIVE 1 (*At*BRI1) forms ligand-independent homo-oligomers, in which the C-terminal domain keeps the kinase in an auto-inhibitory, low basal activity state. Upon Brassinosteroid binding, conformational changes release the kinase from its inhibitory mode to facilitate strong auto-phosphorylation and receptor complex formation with BAK1 (Wang *et al*., 2005). Ligand-independent self-association of *At*FLS2 was reported (Sun *et al*., 2012), but FLIM analysis of *At*FLS2 at the membrane did not show homomerization (Somssich *et al*., 2015). Our FRET-FLIM data show that *At*LORE homomerization does not significantly change within 20 minutes after ligand application (**Fig 1D**). Although receptor complex formation should be rapid, this timeframe was suggested for comparable FLIM studies (Somssich *et al*., 2015). Generally, plants may maintain receptor complexes in a stable, preformed steady-state as a strategy to ensure rapid signaling upon ligand binding that is subsequently fine-tuned by multiple regulators in higher-order complexes.

Our data suggest that homomerization of *At*LORE requires both the ECD and TMD since the ECD or ICD alone barely interact with or outcompete full-length *At*LORE (**Fig 2B, 7C-E**), and SEC shows a monomeric state of *At*LORE-ECDs *in vitro* (**Fig S7D**). The TMD-mediated interaction of *At*LORE-ECD-TMD and *At*LORE-TMD-ICD (**Fig 2D**) supports a significant contribution of the TMD to *At*LORE homomerization. Generally, TMD helixes represent a typical interaction interface in the hydrophobic environment of the membrane and often contain conserved interaction motifs (Langosch & Arkin, 2009; Fink *et al*., 2012; Westerfield & Barrera, 2020), none of which are found in the *At*LORE-TMD. TMDs of Arabidopsis CRINKLY 4 (*At*ACR4) and ACR4-related RLKs have a strong homo-dimerization potential (Stokes & Gururaj Rao, 2008). The *At*ARC4-TMD further mediates the interaction with the RLK CLAVATA1 (Stahl *et al*., 2013). The relevance of TMDs in SD-RLK signaling is underlined by *Os*Pi-d2, in which a SAP in the TMD determines susceptible or resistant alleles towards rice blast (Chen *et al*., 2006). Membrane anchoring is also relevant to *Bra*SRK oligomerization, as velocity sedimentation analytical ultracentrifugation showed that integral SRKs exist as oligomers, while soluble eSRKs and SLGs are primarily monomeric (Giranton *et al*., 2000). We observe a similar behavior for LORE (**Fig 2, Fig S7D**). However, ligand binding seems to work independently of membrane anchoring or dimerization for LORE (**Fig 4, 5**). In *B. rapa*, both SRK:SRK and SRK:SCR interactions contribute to the formation of a tetrameric complex (Ma, R *et al*., 2016; Murase *et al*., 2020). Compared to the large SCR peptide ligands, 3-OH-C10:0 is a small metabolite, which may not be sufficient to crosslink two *At*LORE monomers. One may speculate that LORE dimerization might be rather stabilized by ligand-induced conformational changes, similar to the mechanism described for *At*BRI1 (Wang *et al*., 2005). This is supported by our observation that 3-OH-C10:0 binds to non-dimerizing LORE monomers such as *Alyr*LORE and *Ahal*LORE (**Fig 4C-D, Fig S7**). The molecular mechanism of receptor activation upon ligand binding remains to be elucidated. The partial complementation of *Alyr*LORE in the *lore-*1 phenotype by *Alyr*LORE (**Fig 3**) in *A. thaliana* hints at additional components contributing to LORE receptor complex formation, which might be absent or diversified in *A. lyrata*, like SLG which is absent in the S-locus of *A. lyrata* (Nasrallah, 2023). These components might be recruited upon ligand binding to stabilize LORE homomers and partially rescue *Alyr*LORE homomerization in *A. thaliana*.

*In vitro* phosphorylation assay of microsomal membranes shows that oligomerization of membrane-bound recombinant *Bra*SRKs is essential for auto-phosphorylation (Giranton *et al*., 2000) and that preformed *Bra*SRK oligomers exhibit a basal constitutive auto-phosphorylation. Recombinant cytosolic *At*LORE domains also auto-phosphorylate *in vitro* (Luo *et al*., 2020). We assume that similar to *Bra*SRKs or *At*BRI1, *At*LORE homomerization is required for its phosphorylation, as cell death observed upon *At*LORE overexpression in *N. benthamiana* depends on kinase activity and dimerization (**Fig 1A, 5A**). Pre-formed receptor complexes often seem to have basal constitutive phosphorylation activity that needs to be tightly regulated, as shown for *At*BRI1 (Wang *et al*., 2005). *At*LORE interacts with potentially negatively regulating plant U-box (PUB) E3 ubiquitin ligases (Samuel *et al*., 2008) and the phosphatase LOPP, which attenuates *At*LORE signaling via dephosphorylation (Wang *et al*., 2023). Although LOPP is present in solanaceous tomato (Wang *et al*., 2023) and thus likely also in *N. benthamiana*, further negative regulators of LORE might be absent or limiting in *N. benthamiana*, leading to constant, overshooting signaling and cell death. The phenomenon of cell death-like symptoms upon overexpression of RLKs has been described repeatedly. Overexpression of the (potentially also homomerizing) SD-RLK SPL11 cell-death *s*uppressor 2 (SDS2) in rice or *At*CERK1 in *N. benthamiana* results in spontaneous cell death (Pietraszewska-Bogiel *et al*., 2013; Fan *et al*., 2018). Overproduction of *At*CERK1 or SDS2 might trigger ligand-independent homomerization, leading to auto-activation and dysregulated signaling. *At*LecRK-IX.1 and *At*LecRK-IX.2 exhibit cell death upon overexpression in *A. thaliana* and *N. benthamiana* (Wang *et al*., 2015). Although *At*LecRK-IX.1/2 homomerization was not reported, it is shown for other *At*LecRKs (Guo *et al*., 2018). In *A. thaliana*, overexpression of the co-receptor *At*BAK1, which participates in numerous signaling pathways, results in leaf necrosis and impaired plant development, presumably due to spontaneous activation of multiple receptor complexes (Domínguez-Ferreras *et al*., 2015). Interestingly, all these RLKs, including LORE, are RD-type kinases (Johnson *et al*., 1996) and thus prone to spontaneous auto-activation upon overexpression. In the case of LORE, we exploit the cell death phenotype as a simple read-out for signaling competent receptor complexes regarding kinase activity and homomerization capacity.

Truncated *At*LORE-ECD-TMD exerts a dominant negative effect on 3-OH-C10:0-induced signaling in *N. benthamiana* and *A. thaliana*, while soluble ECD-only versions do not (**Fig 7**). Presumably, this is caused by the formation of signaling incompetent dimers, consisting of a full-length *At*LORE and *At*LORE-ECD-TMD lacking the kinase domain. Competition for ligand binding can be excluded as *Alyr*LORE-ECD-TMD, which binds the ligand to a similar extent but does not homomerize, cannot outcompete *At*LORE signaling. While SLGs were reported to enhance SRK signaling in Brassica (Takayama *et al*., 2001), we do not see a similar phenomenon for the soluble LORE-ECD. However, SLGs show a certain degree of sequence polymorphism compared to their cognate SRK, which might underlie their supportive yet unknown role in establishing SI. Competition-based phenomena have also been described for other RLKs. *Os*LSK1 interacts with itself and five homologous proteins. Overexpression of truncated *Os*LSK1 lacking the kinase domain resulted in increased plant height and yield in rice (Zou *et al*., 2015). Truncated *Os*LSK1 may interfere with signaling of full-length *Os*LSK1 and other interacting SD-RLKs or may compete for ligand binding. Overexpressed *At*FLS2 truncations lacking the LRR domain associate with full-length *At*FLS2 and exert a dominant negative effect on flg22-induced signaling (Sun *et al*., 2012). Expression of a kinase-mutated version of homomerizing *At*BRI1 similarly exerts a dominant effect on Brassinosteroid signaling in a wild-type background (Wang *et al*., 2005).

In conclusion, our study establishes the analysis of dominant negative effects and cell death phenotypes as valuable methods to study LORE homomerization. In addition, natural genetic variation proved to be a powerful tool to elucidate the LORE signalling mechanism. Thus, we were able to show that homomerization of LORE is required for signaling, but apparently in a different way than in *Bra*SRKs, although LORE and *Bra*SRKs share high sequence similarity. Next, it will be interesting to identify the amino acids that cross-link the LORE monomers, decipher the ligand binding site to examine its contribution to receptor activation, and identify further components which may support LORE receptor complex assembly in *A. thaliana*. Overall, our study provides new insights into the activation mechanism of an important model of the large class of SD-RLKs that is widely distributed in plants. Future studies will reveal how widespread the concept of activation through homomerization is within this protein family.

## ACKNOWLEDGEMENTS

We acknowledge the Center for Advanced Light Microscopy (CALM) of imaging@TUM at the TUM School of Life Sciences for providing access to FLIM microscopes, Pascal Falter-Braun (Helmholtz center Munich) and Marcel Quint (MLU Halle-Wittenberg) for providing plant material and Yvonne Stahl (HHU Düsseldorf) for providing FLIM vectors. We thank Julia Santiago and Pedro Jimenez-Sandoval (University of Lausanne) for their help with SEC experiments, and Ralph Hückelhoven and Martin Stegmann for critical discussion of results. Graphs and statistics were created by GraphPad Prism, microscopy images by OMERO and schemes by BioRender.com. Research in the Ranf lab is supported by the German Research Foundation (SFB924/TP-B10 and Emmy Noether program RA-2541/1 to S.R.) and the Swiss National Science Foundation (grant 310030_208139 to S.R.).

## AUTHOR CONTRIBUTIONS

Conception of the project: SR; experimental work and data collection: AF, SEsch, MS, TI, SEib, LJS; data analysis: AF, SEsch, MS, TI, SEib, LJS, SR; data interpretation and discussion of results: SEsch, MS, LJS, SR; drafting the manuscript: SEsch; critical revision of the manuscript: SR; approval of the final manuscript: all authors.

## SUPPORTING INFORMATION

**Figure S1** Phylogenetic tree, multiple sequence alignments and structural models of *At*LORE and other RLKs.

**Figure S2** *At*LORE overexpression causes cell death in *Nicotiana benthamiana*.

**Figure S3** *At*LORE forms homomers in *Nicotiana benthamiana.*

**Figure S4** Localization of *At*LORE truncation variants in *Nicotiana benthamiana*.

**Figure S5** Localization of LORE orthologues in *Nicotiana benthamiana* and ROS response of different *Brassicaceae* species upon flg22 elicitation.

**Figure S6** Protein sequence alignments of cloned *Ahal*LORE sequence compared to the respective genome database sequence.

**Figure S7** Extracellular domains (ECDs) of S-domain receptor like kinases (SD-RLKs) were expressed, harvested and purified from *Nicotiana benthamiana* apoplasts.

**Figure S8** Mapping of dimerization region by analysis of receptor chimera of *At*LORE and *Alyr*LORE in *Nicotiana benthamiana*

**Figure S9** Localization of *At*LORE and *Alyr*LORE domain swaps in *Nicotiana benthamiana*.

**Figure S10** Expression of *At*LORE or *Alyr*LORE truncations which exert a dominant negative effect on full-length *At*LORE signaling in *Nicotiana benthamiana*.

**Figure S11** Expression and localization of *At*LORE-ECD-TMD-mCherry and *At*LORE-ECD-mCherry stably overexpressed in *Arabidopsis thaliana* Col-0^AEQ^.

**Table S1** Protein domain annotations of LORE

**Table S2** Primers used in this study

**Table S3** Annotation of LORE truncations and chimera

**Table S4** List of expression plasmids used in this study

**Table S5** DNA and protein sequence sources

**Table S6** Antibodies used in this study

