## Supplementary Figures for "LORE receptor homomerization is required for 3-hydroxydecanoic acid-induced immune signaling and determines the natural variation of immunosensitivity within the Arabidopsis genus"

A

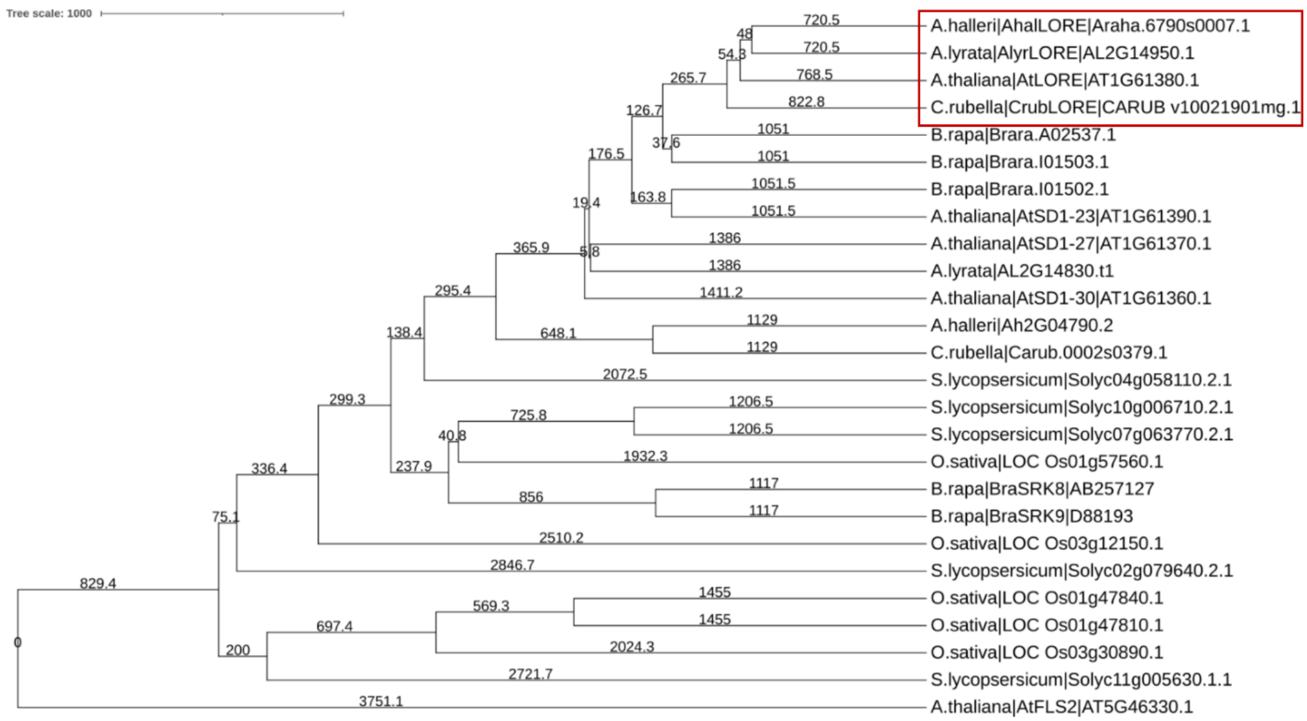

B

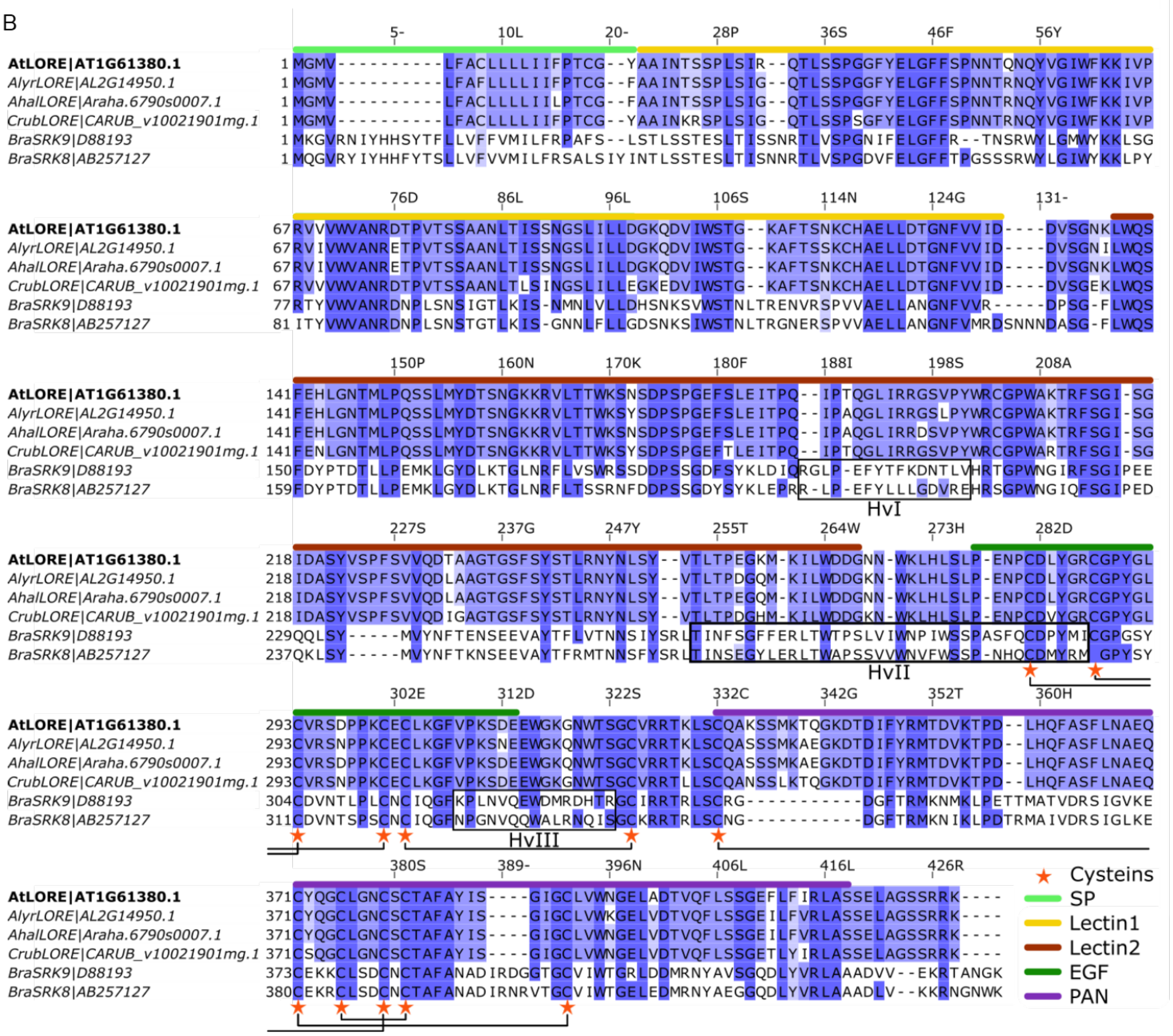
AtLORE|AT1G61380.1
AlyrLORE|AL2G14950.1
AhalLORE|Araha.6790s0007.1
CrubLORE|CARUB_v10021901mg.1
BraSRK9|D88193
BraSRK8|AB257127
HvI
HvII
HvIII
Cysteins
SP
Lectin1
Lectin2
EGF
PAN

C

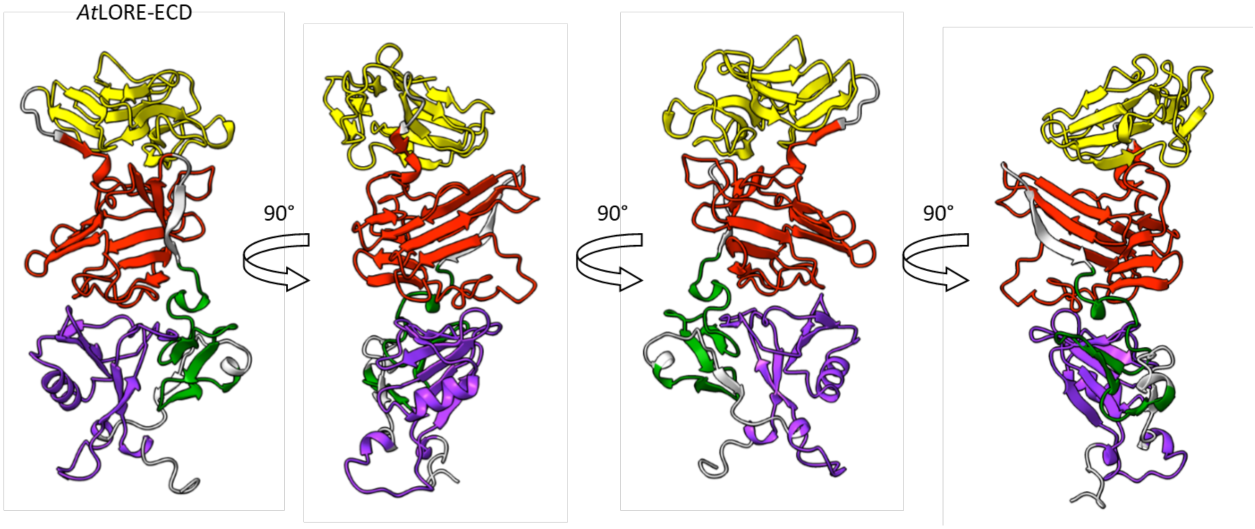
AtLORE-ECD
90°
90°
90°

D

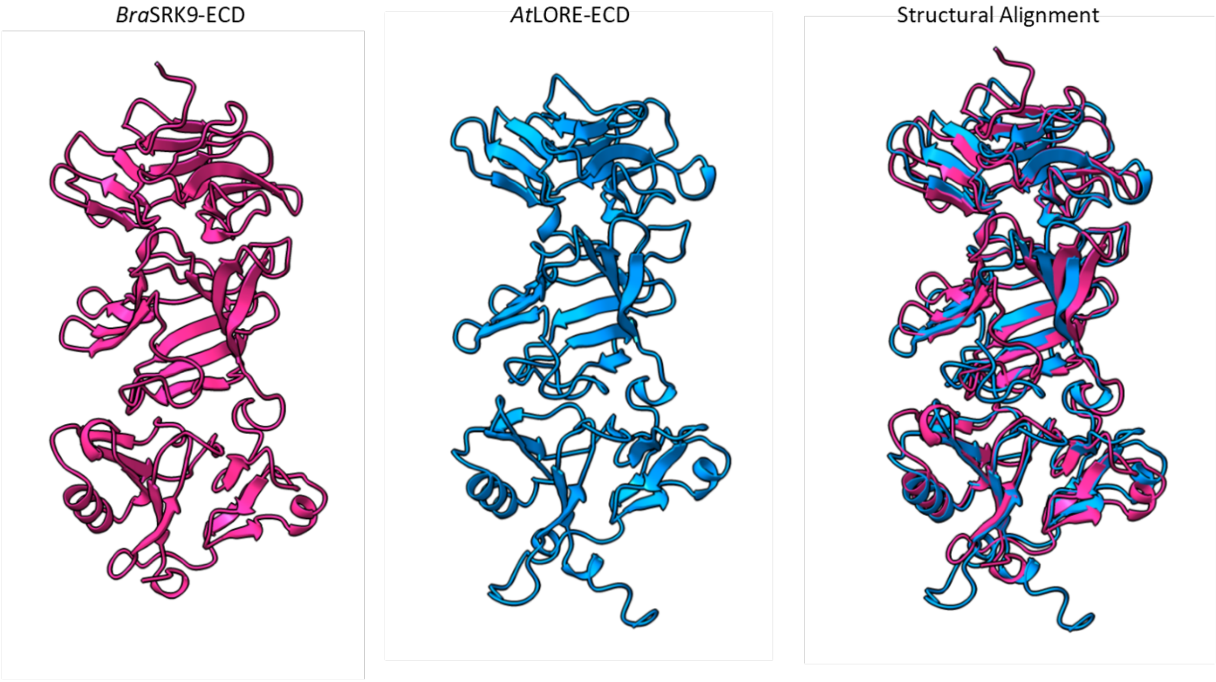
BraSRK9-ECD
AtLORE-ECD
Structural Alignment

E

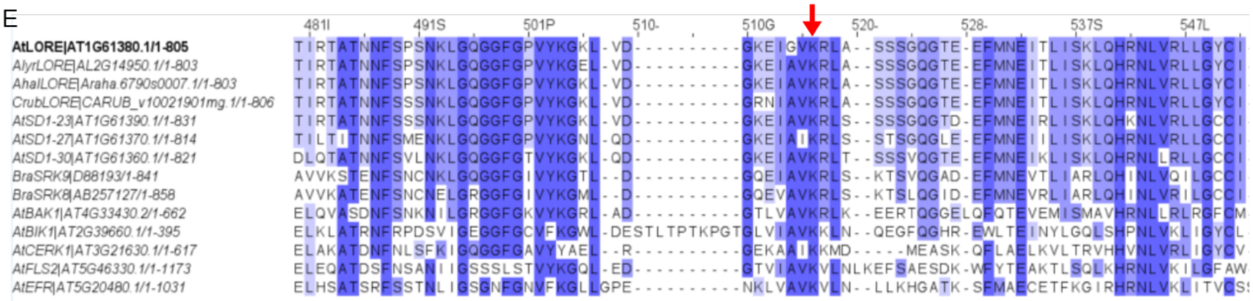
481I
491S
501P
510-
510G
520-
528-
537S
547L
AtLORE|AT1G61380.1/1-805 TIRTATNNFSPSNKLGQGGFGPVYKGKL-VD----------GKEIGVKRLA--SSSGQGTE-EFMNEITLISKLQHRNLVRLLGYCI
AlyrLORE|AL2G14950.1/1-803 TIRTATNNFSPSNKLGQGGFGPVYKGEL-VD----------GKEIAVKRLA--SSSGQGTE-EFMNEITLISKLQHRNLVRLLGYCI
AhalLORE|Araha.6790s0007.1/1-803 TIRTATNNFSPSNKLGQGGFGPVYKGKL-VD----------GKEIAVKRLA--SSSGQGTE-EFMNEITLISKLQHRNLVRLLGYCI
CrubLORE|CARUB_v10021901mg.1/1-806 TIRTATNNFSSSNKLGQGGFGPVYKGKL-VD----------GRNIAVKRLA--SSSGQGTE-EFMNEITLISKLQHRNLVRLLGYCI
AtSD1-23|AT1G61390.1/1-831 TIRTATNNFSSSNKLGQGGFGPVYKGKL-VD----------GKEIAVKRLS--SSSGQGTD-EFMNEIRLISKLQHKNLVRLLGCCI
AtSD1-27|AT1G61370.1/1-814 TILTITNNFSMENKLGQGGFGPVYKGNL-QD----------GKEIAIKRLS--STSGQGLE-EFMNEIILISKLQHRNLVRLLGCCI
AtSD1-30|AT1G61360.1/1-821 DLQTATNNFSVLNKLGQGGFGTVYKGKL-QD----------GKEIAVKRLT--SSSVQGTE-EFMNEIKLISKLQHRNLLRLLGCCI
BraSRK9|D88193/1-841 AVVKSTENFSNCNKLGQGGFGIVYKGTL--D----------GQEIAVKRLS--KTSVQGAD-EFMNEVTLIARLQHINLVQILGCCI
BraSRK8|AB257127/1-858 AVVKATENFSNCNELGRGGFGIVYKGML--D----------GQEVAVKRLS--KTSLQGID-EFMNEVRLIARLQHINLVRILGCCI
AtBAK1|AT4G33430.2/1-662 ELQVASDNFSNKNILGRGGFGKVYKGRL-AD----------GTLVAVKRLK--EERTQGGELQFQTEVEMISMAVHRNLLRLRGFCM
AtBIK1|AT2G39660.1/1-395 ELKLATRNFRPDSVIGEGGFGCVFKGWL-DESTLTPTKPGTGLVIAVKKLN--QEGFQGHR-EWLTEINYLGQLSHPNLVKLIGYCL
AtCERK1|AT3G21630.1/1-617 ELAKATDNFNLSFKIGQGGFGAVYYAEL--R----------GEKAAIKKMD----MEASK-QFLAELKVLTRVHHVNLVRLIGYCV
AtFLS2|AT5G46330.1/1-1173 ELEQATDSFNSANIIGSSSLSTVYKGQL-ED----------GTVIAVKVLNLKEFSAESDK-WFYTEAKTLSQLKHRNLVKILGFAW
AtEFR|AT5G20480.1/1-1031 ELHSATSRFSSTNLIGSGNFGNVFKGLLGPE----------NKLVAVKVLN--LLKHGATK-SFMAECETFKGIRHRNLVKLITVCS

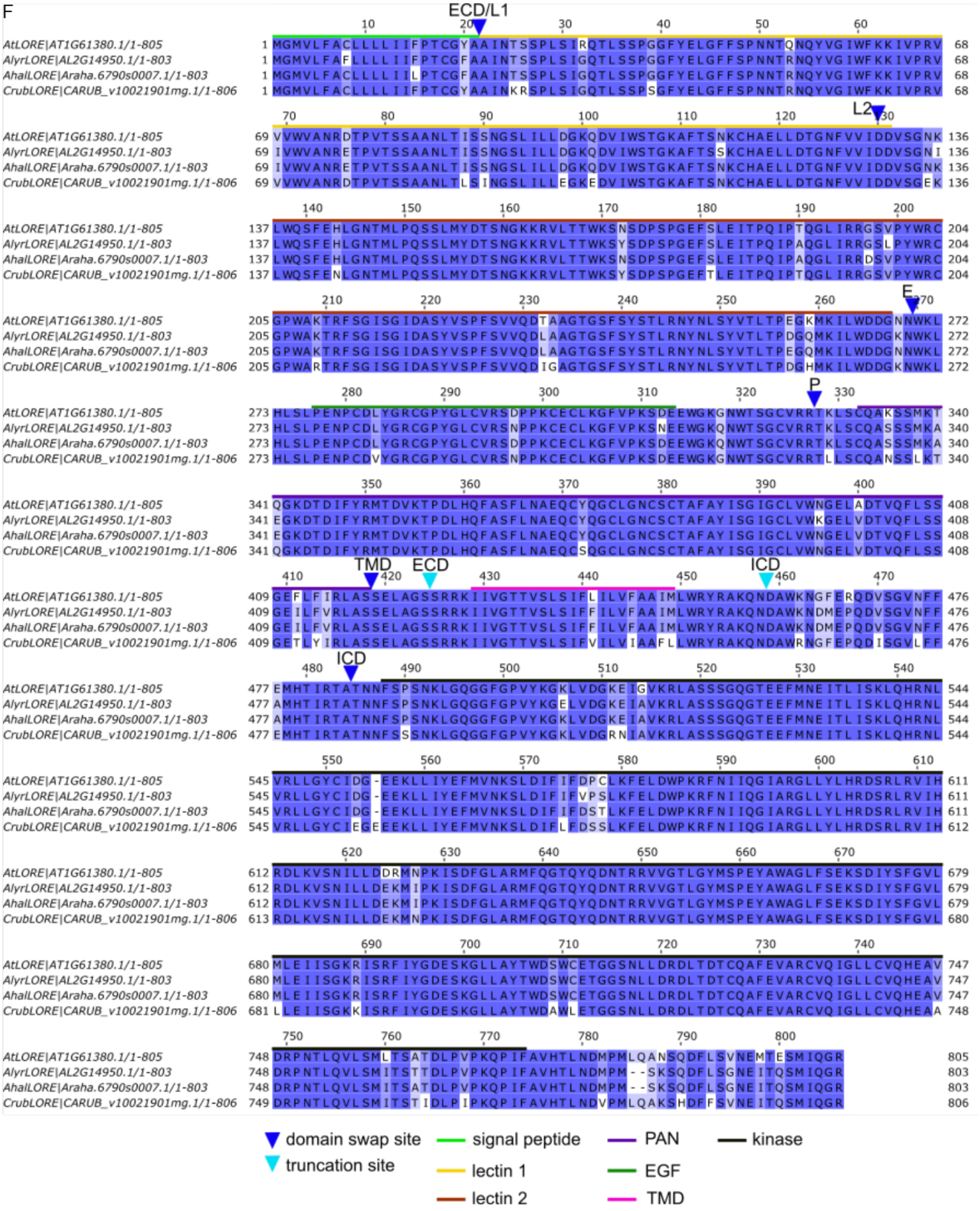
F
ECD/L1
L2
E
P
TMD
ECD
ICD
ICD
AtLORE|AT1G61380.1/1-805
AlyrLORE|AL2G14950.1/1-803
AhalLORE|Araha.6790s0007.1/1-803
CrubLORE|CARUB_v10021901mg.1/1-806
domain swap site
truncation site
signal peptide
lectin 1
lectin 2
PAN
EGF
TMD
kinase

**Figure S1 Phylogenetic tree, multiple sequence alignments and structural models of *At*LORE and other RLKs. A** Multiple protein sequence alignment and phylogenetic analysis of *At*SD-RLKs (*At*SD1-23/27/30/*At*LORE), putative brassicaceous orthologues (top BLAST hits of *At*LORE and *At*SD1-23 against *Arabidopsis halleri, A. lyrata, Capsella rubella* and *Brassica rapa* genomes, Phytozome v13), well-studied *B. rapa* SRKs (*Bra*SRK8/9), SD-RLKs with similar domain architecture as LORE from tomato (*S. lycopersicum*) and rice (*O. sativa*) according to Teixeira *et al.*, 2018 and the LRR-RLK *At*FLS2 (MAFFT alignment, average distance using BLOSUM62, distances are shown, performed with Jalview). The *At*LORE clade is marked with a red box. For detailed sequence source information see supplementary information Table S5. **B** Multiple protein sequence alignment (MAFFT algorithm, Jalview) of the ECDs of *At*LORE, LORE orthologues and well-studied *Bra*SRKs. Blue color intensity indicates percentage identity. *At*LORE was set as reference, and numbering is shown accordingly. LORE protein domain annotations are marked according to Naithani *et al.* (2007) and Uniprot database (Supplementary Information Table S1). Conserved cysteine residues are marked with red stars, and disulfide bonds of *Bra*SRKs are highlighted with brackets. Disulfide bonds and hypervariable regions (Hv) of *Bra*SRKs are marked according to Murase *et al*. (2020). For detailed sequence source information see Supplementary Information Table S5. SP; signal peptide. **C, D** Protein models of the ECDs of *At*LORE (Alphafold model AF-O64782-F1) and *Bra*SRK9 (PDB code: 5GYY; Ma *et al.,* 2016). **C** *At*LORE-ECD protein model with color labeling of protein domains according to **B**. Different side views, rotated by 90° each, are shown. **D** Individual and aligned protein models of *Bra*SRK9-ECD (magenta) and *At*LORE-ECD (blue). Alignment was done with default settings of the Matchmaker tool in ChimeraX (version 1.7.dev202308310222). **E** Protein sequence alignment (MAFFT algorithm) of *At*LORE and other receptor kinases, showing a part of the intracellular kinase domain (*At*LORE T480-I552). Blue color intensity indicates percentage identity. *At*LORE was set as reference, and numbering is shown accordingly. Conserved ATP binding site K516 is marked with a red arrow. **F** Full protein sequence alignment (MAFFT algorithm, Jalview) of *At*LORE and orthologues. Blue color intensity indicates percentage identity. *At*LORE was set as reference, and numbering is shown accordingly Domain annotations, truncation and domain swap sites are labelled as indicated. Detailed domain annotations are given in Supplementary Tables S1 and S3.

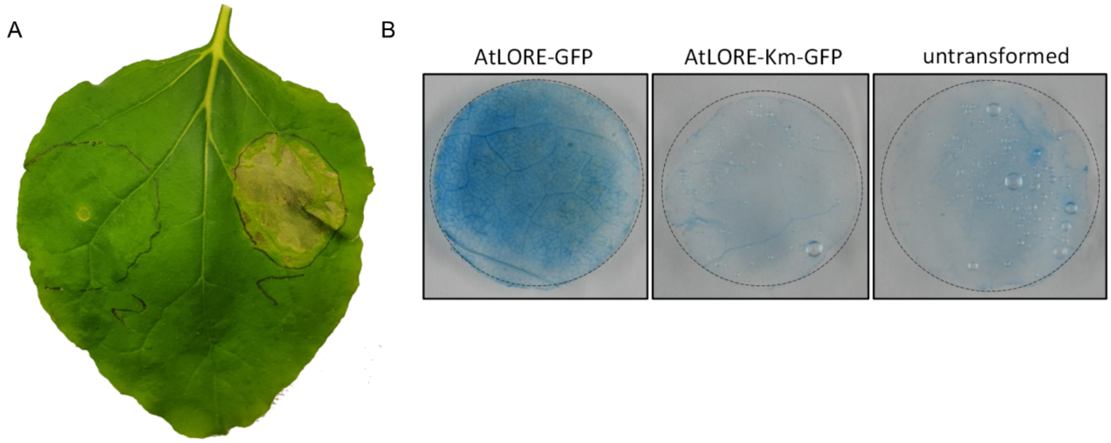


**Figure S2 *At*LORE overexpression causes cell death in *Nicotiana benthamiana*.** **A** Macroscopic cell death symptoms of *N. benthamiana* overexpressing (CaMV35S promoter) *At*LORE-Km-GFP (left, number 2) or *At*LORE-GFP (right, number 1) six days post agro-infiltration. Infiltrated area is marked with black circles. **B** Trypan blue staining of leaf discs (Ø 2 cm) from *N. benthamiana* transiently overexpressing (CaMV35S promoter) *At*LORE or *At*LORE-Km five days post agro-infiltration or an untransformed control.

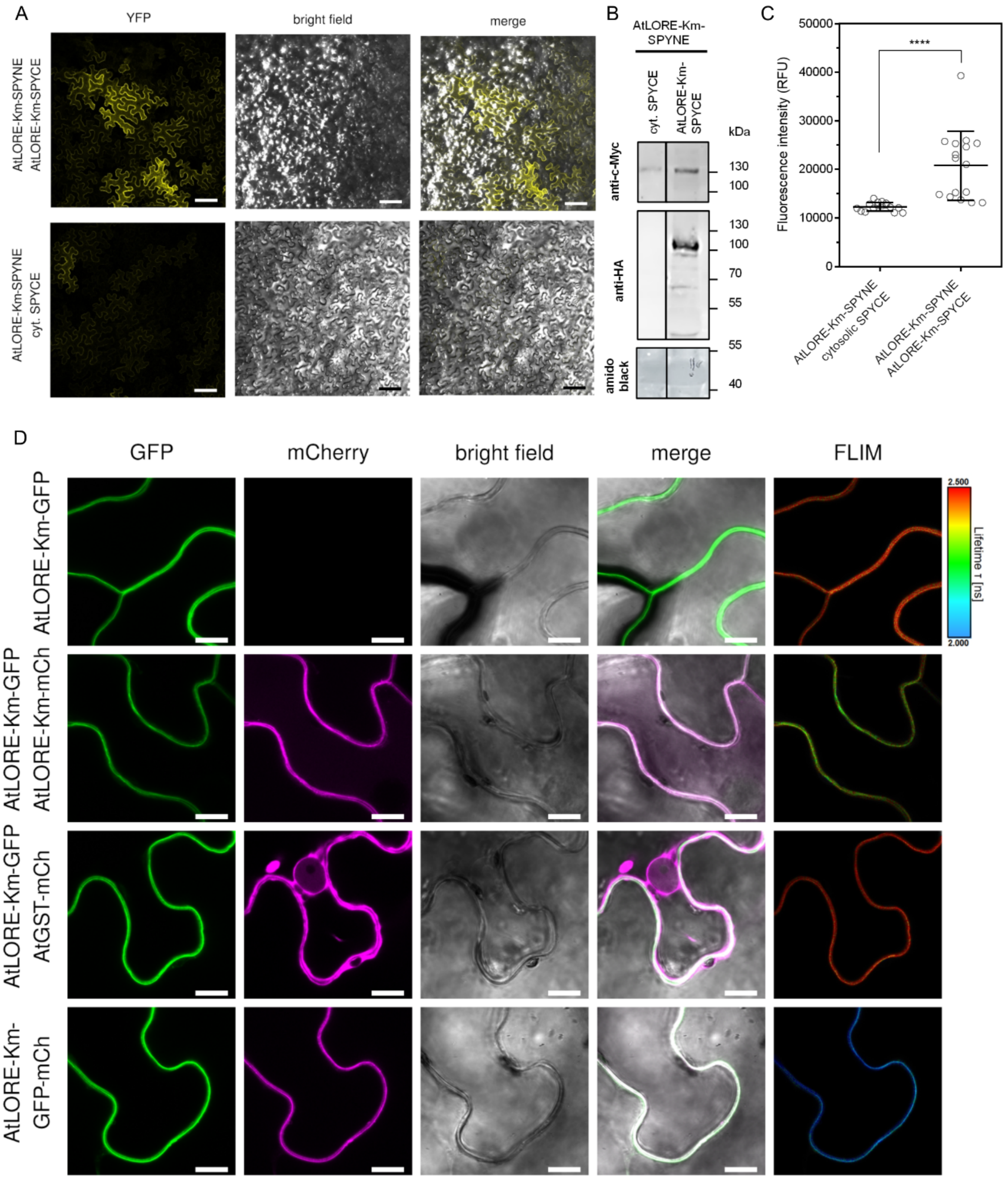


**Figure S3 *At*LORE forms homomers in *Nicotiana benthamiana.*** **A** Confocal laser scanning microscopy of Bimolecular Fluorescence complementation (BiFC) experiments two days post agro-infiltration. SPYNE- and SPYCE-tagged interaction candidates were transiently expressed (CaMV35S promoter) in *N. benthamiana*. Z-stack of 22 Z-sections is shown. Scale bar represents 100 µm. **B** Anti-c-Myc and anti-HA immunoblot of proteins of BiFC experiment. SPYCE includes a c-Myc tag, SPYNE an HA tag. Total protein was stained with amido black. Cytosolic SPYCE was too small to resolve in this blot. **C** Fluorescence intensity quantification of BiFC experiment. Data show mean with SD; n=16 leaf discs. Statistics were analyzed by unpaired t-test; ****, P<0.0001. **D** Confocal laser scanning microscopy and FLIM images show subcellular localization of proteins used for FRET-FLIM analysis (Fig. 1 C, D) transiently expressed in *N. benthamiana.* Scale bar represents 10 µm. Rainbow-colorscale of FLIM images represents mean GFP lifetime (arrivel time of photons after laser pulse) ranging from 2.0 to 2.5 ns.

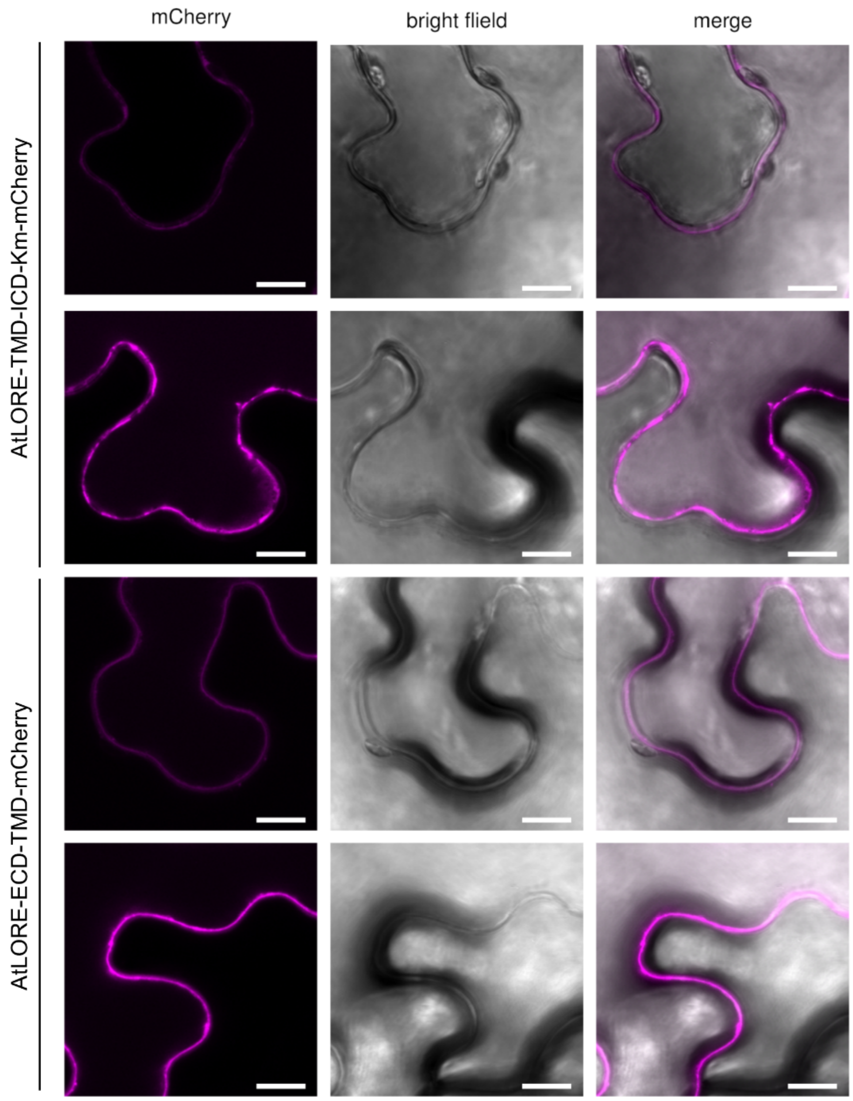


**Figure S4 Localization of *At*LORE truncation variants in *Nicotiana benthamiana***. Confocal laser scanning microscopy shows subcellular localization of the indicated *At*LORE-Km-mCherry truncations used for FRET-FLIM analysis (Fig. 2 C, D) transiently expressed (estradiol-inducible XVE promotor) in *N. benthamiana.* Two representative images with moderate and strong expression are shown. Scale bar represents 10 µm.

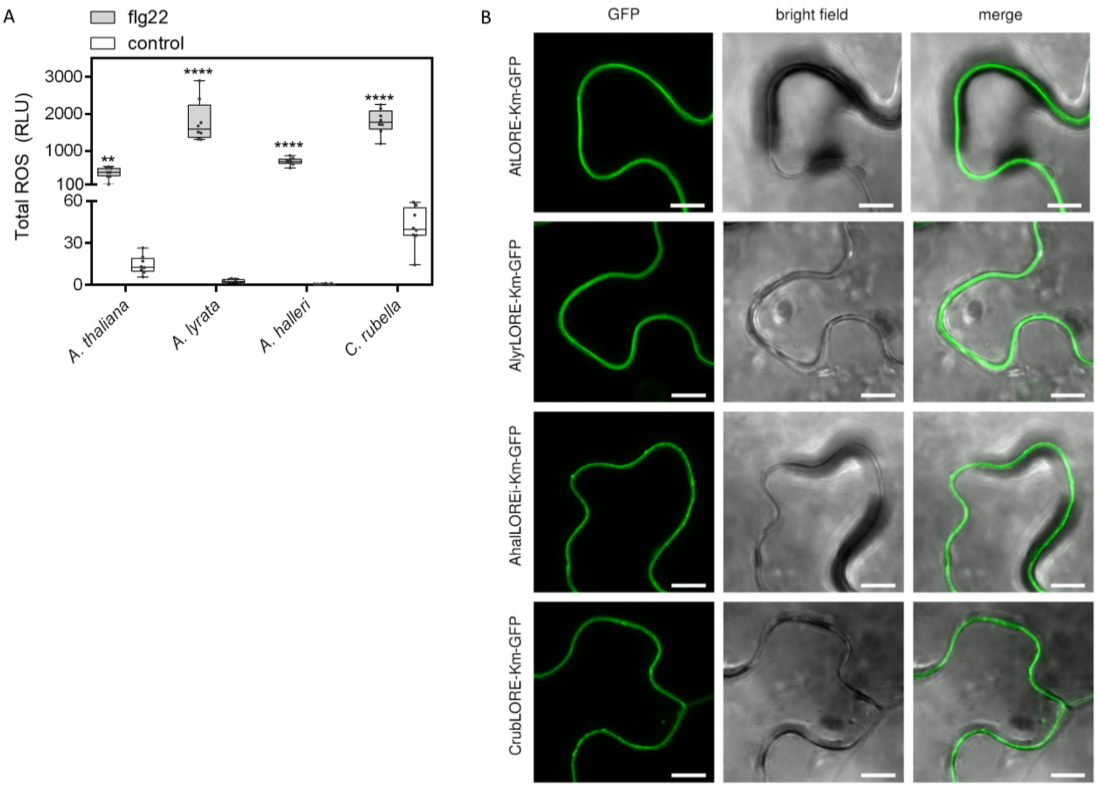


**Figure S5 Localization of LORE orthologues in *Nicotiana benthamiana* and ROS response of different *Brassicaceae* species upon flg22 elicitation. A** Total ROS accumulation of *A. halleri*, *C. rubella*, *A. lyrata* and *A. thaliana* leaf discs elicited with 500 nM flg22. Control represents MeOH treatment identical to data in Figure 3A. Median with minimum to maximum of total ROS of 3-60 minutes after elicitation is shown (n=8 leaf discs). Statistics were analyzed by two-way ANOVA with Sidak's multiple comparisons test, ****, $P<0.0001$; **, $P\leq0.0021$. **B** Localization of GFP-tagged LORE orthologues was analyzed by confocal laser scanning microscopy upon transient expression (estradiol-inducible XVE promotor) in *N. benthamiana.* Scale bar represents 10 µm.

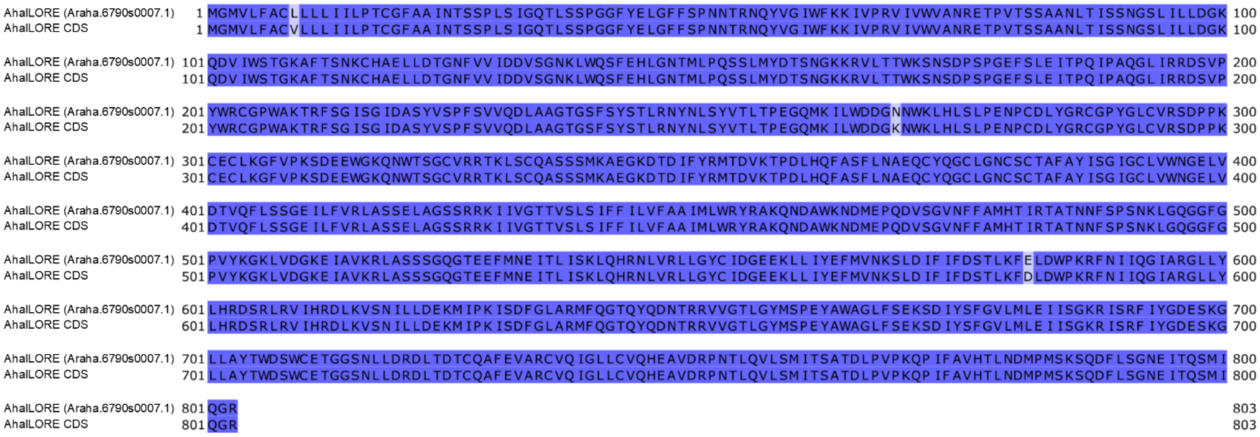


**Figure S6 Protein sequence alignments of cloned *Ahal*LORE sequence compared to the respective genome database sequence.** Amino acid sequence of Araha.6790s0007.1 was obtained from Phytozome v13 (https://phytozome-next.jgi.doe.gov) and compared to the sequence of *Ahal*LORE CDS cloned from cDNA. Blue color intensity indicates sequence identity. Alignment was performed with Jalview (MAFFT algorithm).

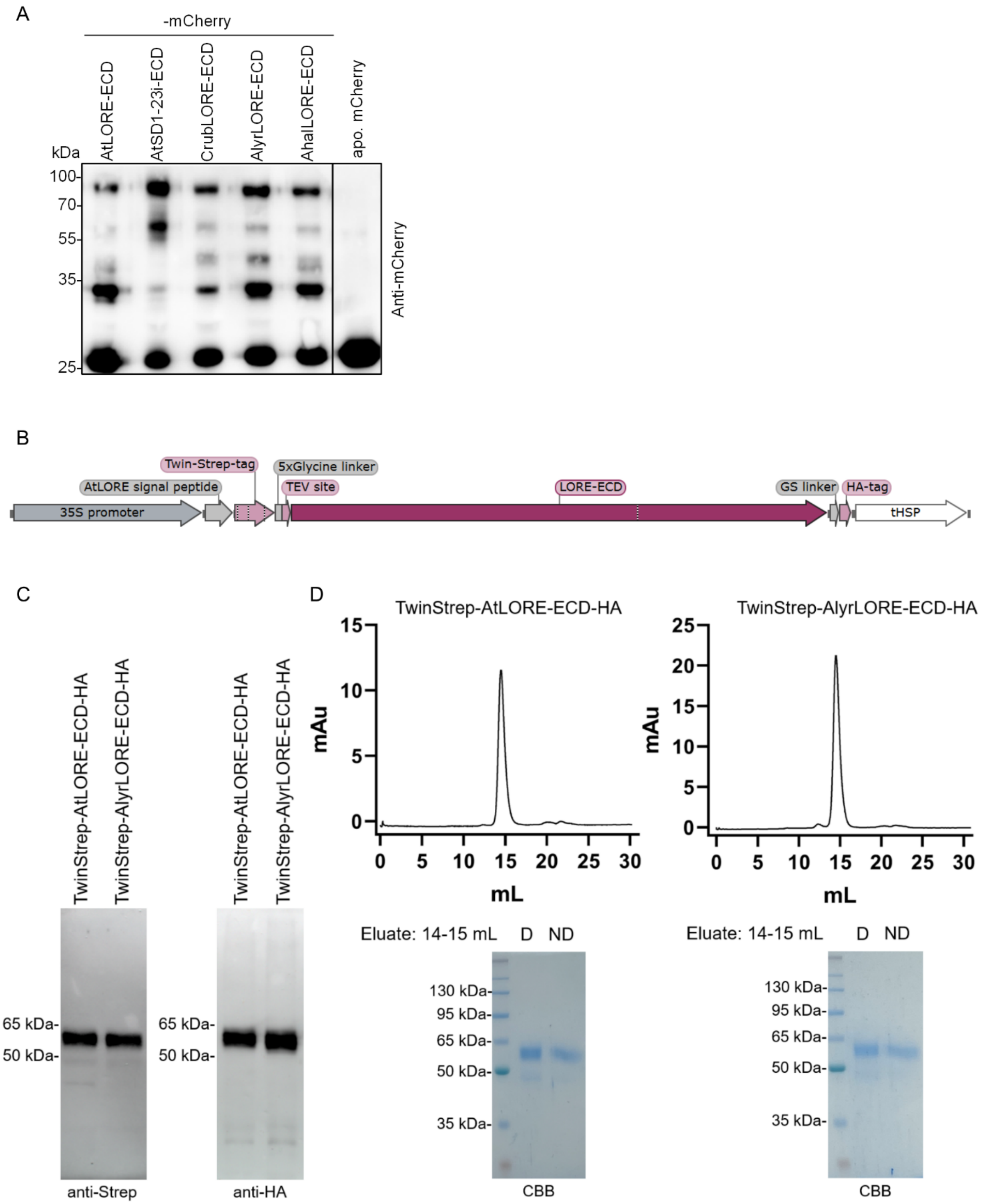


**Figure S7 Extracellular domains (ECDs) of S-domain receptor-like kinases (SD-RLKs) were expressed, harvested and purified from *Nicotiana benthamiana* apoplasts. A** Anti-mCherry immunoblot of apoplastic washing fluids (AWF) of *N. benthamiana* expressing (CaMV35S promoter) mCherry-tagged SD-RLK ECDs or apoplastic (apo.) mCherry. 5 µL of concentrated

AWF with a total protein concentration of 1.5 mg/mL was loaded for each sample. **B** Scheme of expression construct of LORE-ECD with N-terminal Twin-strep-tag and C-terminal HA-tag for protein purification. **C** Anti-Strep-tag (left) and anti-HA (right) immunoblot of *At*LORE-ECD and *Alyr*LORE-ECD from apoplastic washing fluids of *N. benthamiana*. **D** Size exclusion chromatography (SEC) of *At*LORE-ECD and *Alyr*LORE-ECD. The 14$^{th}$ to 15$^{th}$ mL fractions of SEC analysis (upper panel) were subjected to SDS-PAGE analysis with denaturing and non-denaturing treatments (lower panel). D, Laemmli sample buffer at 95$^{o}$C; ND, non-reducing LDS buffer; CBB, Coomassie Brilliant Blue staining.

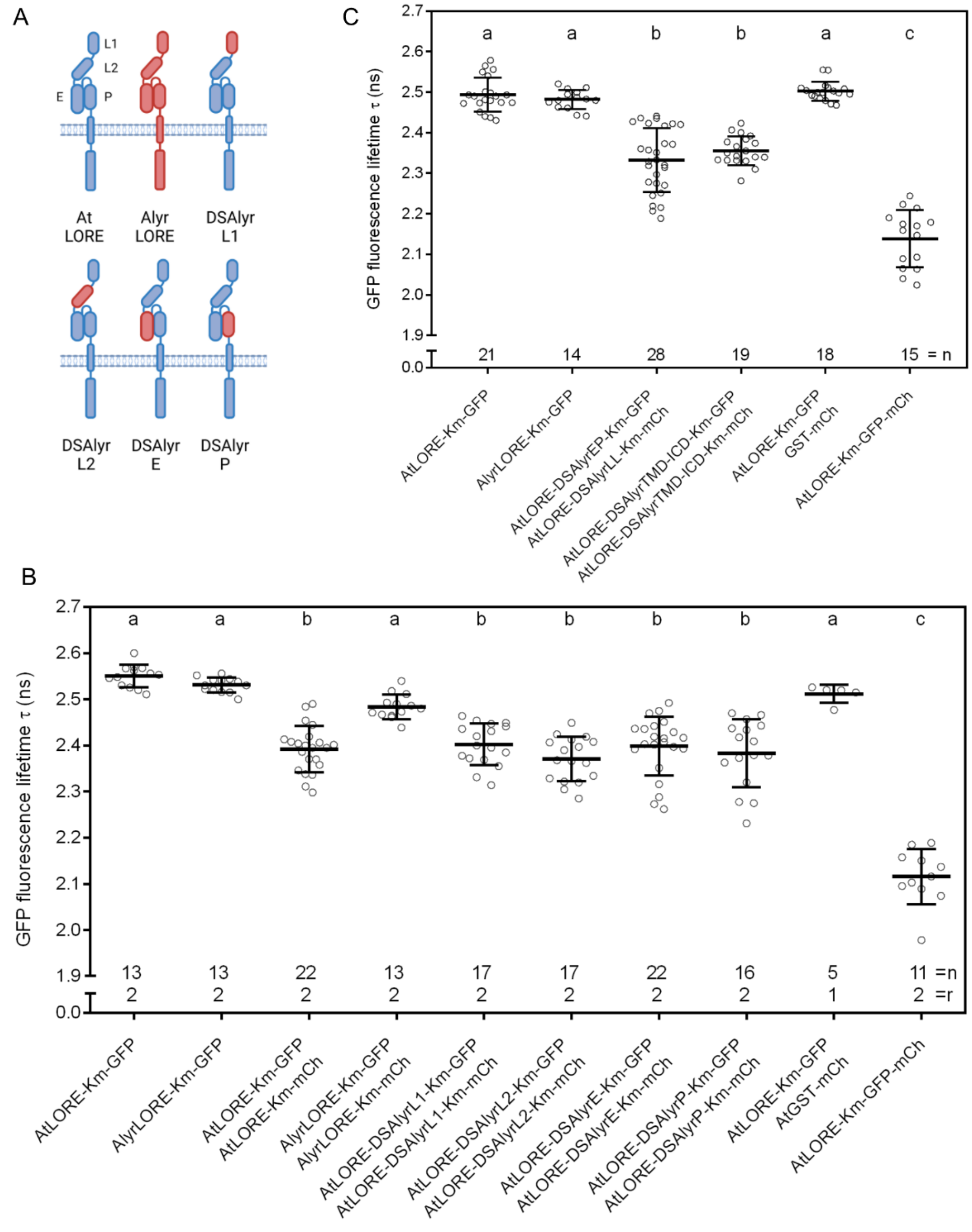


**Figure S8 Mapping of dimerization region by analysis of receptor chimera of *At*LORE and *Alyr*LORE in *Nicotiana benthamiana*. A** Schematic overview of chimera between single domains of *Alyr*LORE and *At*LORE. DS, domain swap; L1, lectin domain 1; L2, lectin domain 2; E, EGF domain, P, PAN domain. **B, C** FRET-FLIM analysis of different chimera between *At*LORE and *Alyr*LORE transiently expressed (estradiol-inducible XVE promotor) in *N. benthamiana.* Data show mean with SD of pooled data from several biological replicates (B: r=2, C: as indicated). n, number of analyzed cells; r, number of biological replicates; mCh, mCherry. Statistics were analyzed by one-way ANOVA with Tukey‘s multiple comparisons test, α=0.01. Data not sharing the same letter are significantly different.

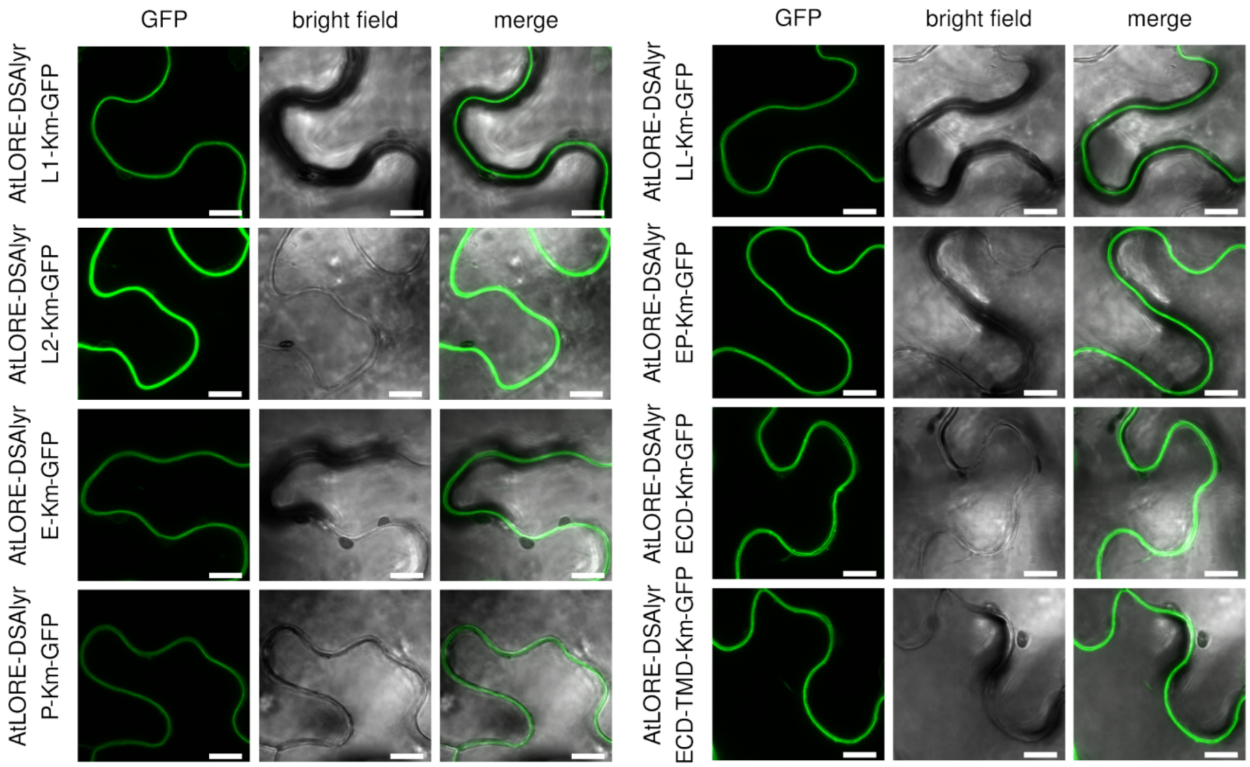


**Figure S9 Localization of *At*LORE and *Alyr*LORE domain swaps in *Nicotiana benthamiana*.** Localization of GFP-tagged proteins was analyzed by confocal laser scanning microscopy upon transient expression (estradiol-inducible XVE promotor) in *N. benthamiana.* Scale bar represents 10 µm.

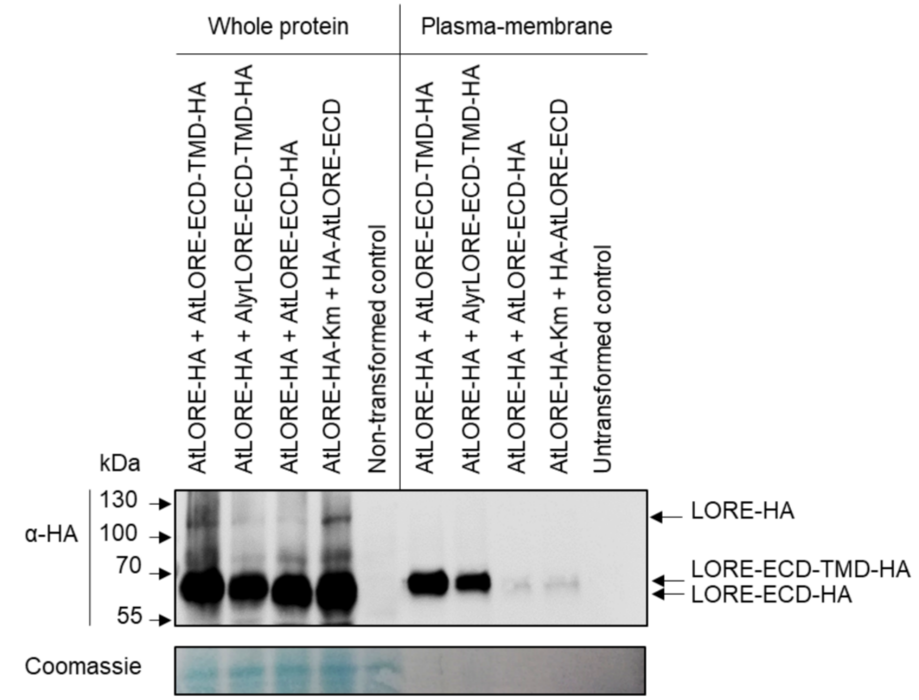


**Figure S10 Expression of *At*LORE or *Alyr*LORE truncations that exert a dominant negative effect on full-length *At*LORE signaling in *Nicotiana benthamiana*.** Anti-HA immunoblot of *At*LORE or *At*LORE-Km transiently coexpressed (CaMV35S promoter) with *At*LORE or *Alyr*LORE truncation variants (LORE-ECD-TMD-HA; HA-LORE-ECD) in *N. benthamiana*. Whole protein and enriched plasma membrane fractions were collected by Minute™ Plant Plasma Membrane Protein Isolation Kit (Invent Biotechnologies, Inc., SM-005). Total protein was stained with Coomassie brilliant blue.

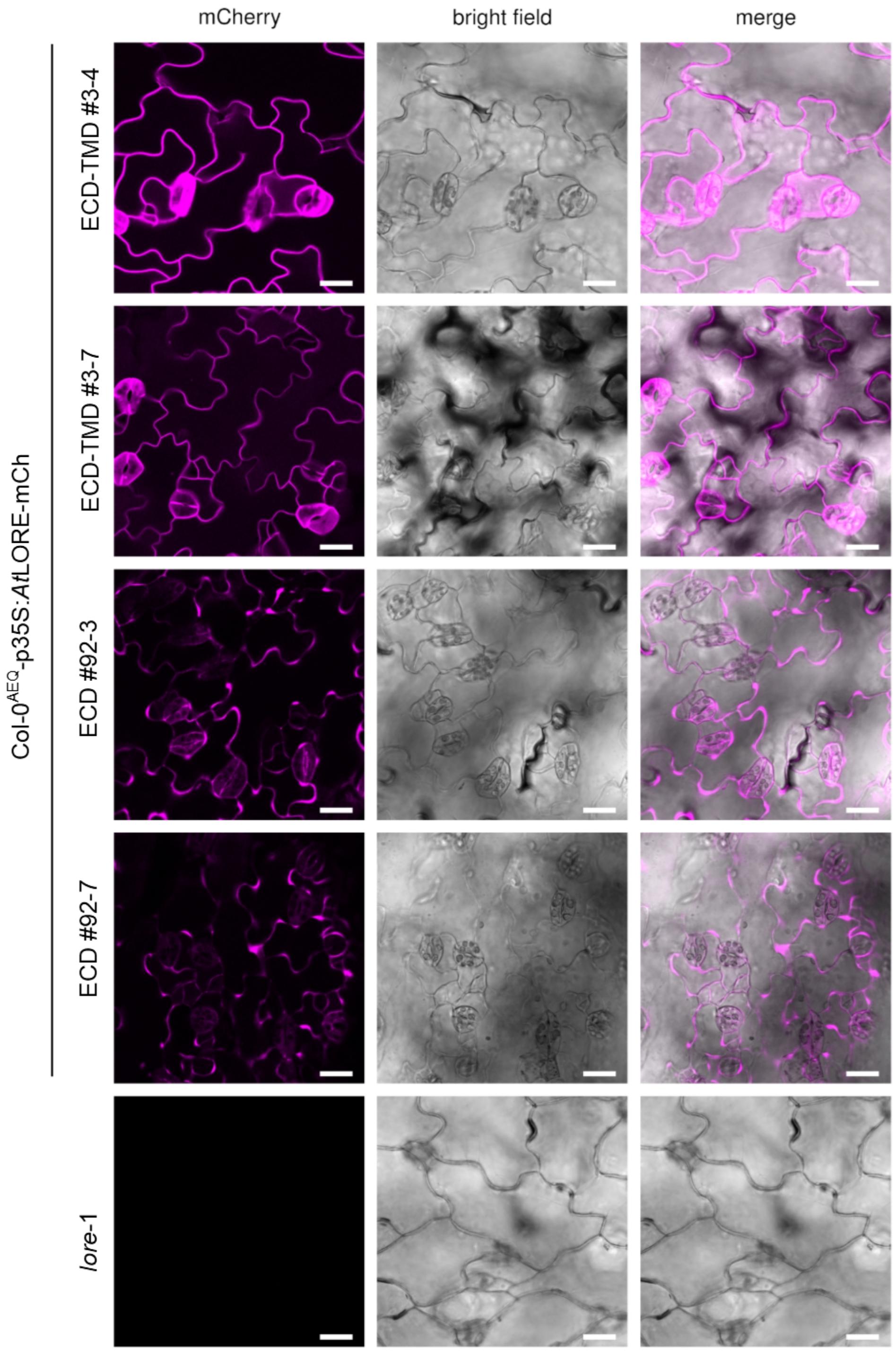


**Figure S11 Expression and localization of *At*LORE-ECD-TMD-mCherry and *At*LORE-ECD-mCherry stably overexpressed in *Arabidopsis thaliana* Col-0$^{AEQ}$.** FastRed selected seedlings of the segregating T2 generation of two independent transgenic lines were grown in liquid culture and mCherry signals were analyzed via confocal laser-scanning microscopy; scale bar represents 20 µm.
