## Supplementary Tables for "LORE receptor homomerization is required for 3-hydroxydecanoic acid-induced immune signaling and determines the natural variation of immunosensitivity within the Arabidopsis genus"

**Table S1 *At* LORE domain annotations**

*At* LORE domain annotations according to Uniprot O64782 and adapted from *Bra* SRKs in Naithani *et al.* , 2007

| Domain | Amino acids | Source |
| --- | --- | --- |
| Signal peptide | 1-21 | Uniprot: O64782 |
| Lectin domain 1 | 22-131 | Naithani *et al.* , 2007 |
| Lectin domain 2 | 137-267 | Naithani *et al* ., 2007 |
| EGF domain | 277-313 | Uniprot: O64782 |
| PAN domain | 332-418 | Uniprot: O64782 |
| Transmembrane domain | 429-449 | Uniprot: O64782 |
| Kinase domain | 488-773 | Uniprot: O64782 |
| ATP binding site | K516 | Uniprot: O64782; Luo *et al* ., 2020 |

**Table S2 Primer**

**CDS cloning from cDNA or vectors**

| Construct name | Source | primer name | sequence 5' to 3' |
|---|---|---|---|
| *Alyr* LORE | AL2G04470, *A. lyrata cDNA* | Alyr-START | tgaagacttAATGGGTATGGTTTTATTTGC |
| | | Alyr-STOPm | tgaagacttACTAGCGTCCTTGGATCATAGATTG |
| *Ahal* LORE | Araha.6790s0007, *A.halleri* cDNA | Ahal-START | ttgaagacttAATGGGTATGGTTTTATTTGC |
| | | Ahal-STOP | ttgaagacttACTATCTTCCTTGAATCATAG |
| gluthation transferase (*At* GST) | AT1G17170, amplified from pGEX-6P-1 | START-GST_F | tgaagacttaATGTCCCCTATACTAGGTTATTGG |
| | | STOP-GST_R | tgaagacttactaTTTTGGAGGATGGTCGCCAC |
| cytosolic GFP | GFP | GFP-Ntag_F | tgaagacttaATGGTGAGCAAGGGC |
| | | GFP-STOP | tgaagacttactATTTGTACAGCTCGTCCATG |
| cytosolic mCherry | mCherry | mCherry-Ctag_F | tgaagactttCCCGGGATGGTGAGCAAGGGC |
| | | mCherry-BpiM_R | tGAAGACcgTtTTCTTCTGCATTACGG |
| | | mCherry-BpiM_F | tGAAGACagAAaACGATGGGCTGG |
| | | mCherry-Ctag_R | tgaagacttaCTATTTGTACAGCTCGTCCATG |
| *At* LORE | Ranf et al., 2015 | - | - |
| *At* LORE-Km | Ranf et al., 2015 | | |
| *Crub* LORE | Shu et al., 2021 | - | - |
| *At* SD1-23i-ECD | Shu et al., 2021 | - | - |
| apoplastic mCherry | Shu et al., 2021 | - | - |

**Intron cloning *Ahal* LORE**

| Construct name | Target | primer name | sequence 5' to 3' |
|---|---|---|---|
| *Ahal* LOREi | Araha.6790s0007, *AhalLORE* cDNA | Ahal-Intron-F | tgaagacttagcttcttcagtcc |
| | | Ahal-Intron-R | tgaagacttaccctaactcataaaatcc |
| SD1-29-intron | At1g61380, *At* LORE gDNA | 129intron-F | tttgaagacttGGgtaaggataaaatacatttcttcc |
| | | 129intron-R | tttgaagacgaAGCTGAGATATTCACCAG |

**Site directed mutagenisis (Km-versions)**

| Construct name | Target | primer name | sequence 5' to 3' |
|---|---|---|---|
| *Ahal* LORE-Km | *Ahal* LORE-CDS | Ahal-K516A-F | gaaatagctgttgcacgccttgctagtagc |
| | | Ahal-K516A-R | gctactagcaaggcgtgcaacagctatttcc |
| *Alyr* LORE-Km | *Alyr* LORE-CDS | Alyr-K516A-F | tGAAGACttTgcaCGCCTTGCTAGTAGCTCC |
| | | Alyr-K516A-R | tGAAGACttTgcaACAGCTATTTCCTTCCCATC |
| *Crub* LORE-Km | *Crub* LORE-CDS | Crub_K516A_QC_f | GGGAGGAATATAGCTGTTGCGCGCCTTGCTAGTAGCTCCG |
| | | Crub_K516A_QC_r | CGGAGCTACTAGCAAGGCGCGCAACAGCTATATTCCTCCC |

**Cloning of *At* LORE truncations**

| Construct name | Target | primer name | sequence 5' to 3' |
|---|---|---|---|
| *At* LORE-TMD-ICD | *At* LORE CDS in GoldeGate entry vector | SD129-TM-F | tgaagacttAGCTGGAAGCAGTCGAAGG |
| | | SD129-SP-R | tgaagacttAGCTGCATAGCCACAAG |
| *At* LORE-ECD | *At* LORE CDS in GoldeGate entry vector | SD129-START-Bpi | tgaagacttAATGGGTATGGTTTTATTTGCTTGC |
| | | SD129-EC-R | tgaagacttACTAGCTTCCAGCCAATTCTG |
| *At* LORE-ECD-TMD | *At* LORE CDS in GoldeGate entry vector | SD129-START-Bpi | tgaagacttAATGGGTATGGTTTTATTTGCTTGC |
| | | SD129-EC-TM-R | tgaagacttACTAATTTTGTTTCGCTCTGTATC |
| *At* LORE-ICD | *At* LORE CDS in GoldeGate entry vector | SD129-ICD-F | tgaagacttaatgGATGCATGGAAGAATGG |
| | | DS-START-EP21-R | tgaagacCCCATttgagacgatataCTGC |
| *Alyr* LORE-ECD | AlyrLORE CDS in GoldenGate entry vector | AlyrLORE-START-BpiI-F | tgaagacttaatgggtatggttttatttgc |
| | | AlyrLORE-EC-BpiI_R | tgaagacttaCTAGCTTCCAGCCAATTCTG |
| *Alyr* LORE-ECD-TMD | AlyrLORE CDS in GoldenGate entry vector | SD129-EC-TM_R | tgaagacttaCTAATTTTGTTTCGCTCTGTATC |
| | | pGGEntL-BB-STOP_F | GCGGCGGAAGACTTTAGtagagacgTCCGCCTCC |
| AhalLORE-ECD | AhalLORE CDS in GoldenGate entry vector | AhalLORE-START-BpiI_F | tttgaagacttaatgggtatggttttatttgc |
| | | SD129-EC-R-BpiI_R | TGAAGACTTACTAGCTTCCAGCCAATTCTG |

**Cloning *At* LORE-*Alyr* LORE chimera**

| Construct name | AtLORE-region | primer name | sequence 5' to 3' |
|---|---|---|---|
| *At* LORE-AlyrDS-ECD | AtTMD-ICD-backbone | SD129-DSN-F | tgaagacAGTTCAGAATTGGCTGGAAG |
| | | DS-START-EP21-R | tgaagacCCCATttgagacgatataCTGC |
| | AlyrECD | Alyr-START | tgaagacttaatgggtatggttttatttgc |
| | | ALYR-DSN-R | tgaagactcTGAACTTGCAAGACGAAC |
| *At* LORE-AlyrDS-ECD-TMD | AtICD -backbone | SD129-DSP-F | tgaagacgcCACCAATAACTTCAGTCC |
| | | DS-START-EP21-R | tgaagacCCCATttgagacgatataCTGC |
| | AlyrECD-TMD | Alyr-START | tgaagacttaatgggtatggttttatttgc |
| | | ALYR-DSP-R | tgaagacttGGTGGCAGTTCGTATG |
| *At* LORE-AlyrDS-ICD | AtECD-TMD-backbone | DS-STOP-EP21-F | tgaagacggTAGTAGAGACGTCCGC |
| | | SD129-DSN-R | tgaagactcTGAACTTGCAAGACGAATG |
| | AlyrICD | SD129-DSP-F | tgaagacgcCACCAATAACTTCAGTCC |
| | | Alyr-STOPm | tgaagacttACTAGCGTCCTTGGATCATAGATTG |
| *At* LORE-AlyrDS-TMD-ICD | AtECD-backbone | DS-STOP-EP21-F | tgaagacggTAGTAGAGACGTCCGC |
| | | SD129-DSP-R | tgaagacttGGTGGCAGTTCGTATTG |
| | AlyrTMD-ICD | ALYR129-DSN-F | tgaagacagTTCAGAATTGGCTGGAAG |
| | | Alyr-STOPm | tgaagacttACTAGCGTCCTTGGATCATAGATTG |
| *At* LORE-AlyrDS-LL | AtEP-TMD-ICD-backbone | SD129_DSA_R | tttgaagacttTATAGCTGCATAGCCACAAG |
| | | SD129_DSC_F | tttgaagacaaTTGGAAGCTTCACTTGTC |

| Construct name | AtLORE-region | primer name | sequence 5' to 3' |
|---|---|---|---|
| | AlyrLL | ALYR129_DSA_F | tttgaagacgcTATAAACACATCAAGTCc |
| | | ALYR129_DSC_R | ttgaagacttCCAATTTTTTCCATCATCCC |
| *At* LORE-AlyrDS-EP | AtLL-TMD-ICD-backbone | SD129_DSC_R | tttgaagacttCCAATTATTTCCATCATCCC |
| | | SD129-DSN-F | tgaagacagTTCAGAATTGGCTGGAAG |
| | AlyrEP | ALYR129_DSC_F | tttgaagacaaTTGGAAGCTTCACTTGTC |
| | | ALYR129-DSN-R | tgaagactcTGAACTTGCAAGACGAAC |
| *At* LORE-AlyrDS-L1 | AtL2-EP-TMD-ICD-backbone | SD129_DSA_R | tttgaagacttTATAGCTGCATAGCCACAAG |
| | | SD129_DSB_F | tttgaagacatTGATGATGTTTCAGGG |
| | AlyrL1 | ALYR129_DSA_F | TTTGAAGACGCtataaacacatcaagtc |
| | | ALYR129_DSB_R | TTTGAAGACTCatcaattacaacaaaatttc |
| *At* LORE-AlyrDS-L2 | AtL1-EP-TMD-ICD-backbone | SD129_DSB_R | tttgaagactcATCAATTACAACAAAATTTC |
| | | SD129_DSC_F | tttgaagacaaTTGGAAGCTTCACTTGTC |
| | AlyrL2 | ALYR129_DSB_F | TTTGAAGACATtgatgatgtttcaggg |
| | | ALYR129_DSC_R | TTTGAAGACTTccaattttttccatcatc |
| *At* LORE-AlyrDS-E | AtLL-P-TMD-ICD-backbone | SD129_DSC_R | tttgaagacttCCAATTATTTCCATCATCCC |
| | | SD129_DSD_F | tttgaagaccgTACAAAATTATCTTGCCAAG |
| | AlyrE | ALYR129_DSC_F | TTTGAAGACAAttggaagcttcacttgtc |
| | | ALYR129_DSD_R | TTTGAAGACTTgtacgcctcacacaccc |
| *At* LORE-AlyrDS-P | AtLL-E-TMD-ICD-backbone | SD129_DSE_R | tttgaagacttTGTACGTCTaACACACCCAC |
| | | SD129-DSN-F | tgaagacagTTCAGAATTGGCTGGAAG |
| | AlyrP | ALYR129_DSE_F | TTTGAAGACCGtacaaaattatcttgccaggc |
| | | ALYR_DSN_Bpi1_R | GAAGACTTtgaacttgcaagacgaacg |

**Table S3 Annotation of LORE truncations and chimera**

**LORE truncations**

| Truncation construct | Amino acids present | Amino acids deleted |
|---|---|---|
| LORE-ECD | 1-424 | 425-805 |
| LORE-ECD-TMD | 1-458 | 459-805 |
| LORE-TMD-ICD | 1-21, 422-805 | 22-421 |
| LORE-ICD | 459-805 | 1-458 |

***At* LORE-*Alyr* LORE chimera/domain swaps**

| | Amino Acids | |
|---|---|---|
| Domain Swap construct | *At* LORE-region | *Alyr* LORE-region |
| *At* LORE-DS*Alyr* ECD | 419-805 | 1-418 |
| *At* LORE-DS*Alyr* ECD-TMD | 485-805 | 1-484 |
| *At* LORE-DS*Alyr* ICD | 1-484 | 485-805 |
| *At* LORE-DS*Alyr* -TMD-ICD | 1-418 | 419-805 |
| *At* LORE-DS*Alyr* -LL | 1-22, 270-805 | 23-269 |
| *At* LORE-DS*Alyr* -EP | 1-269, 419-805 | 270-418 |
| *At* LORE-DS*Alyr* -L1 | 1-22, 130-805 | 23-129 |
| *At* LORE-DS*Alyr* -L2 | 1-129, 270-805 | 130-269 |
| *At* LORE-DS*Alyr* -E | 1-269, 328-803 | 270-327 |
| *At* LORE-DS*Alyr* -P | 1-327, 419-805 | 328-418 |

**Table S4 List of expression plasmids**

For Co-IP, ROS, Chlorophyll fluorescence, BiFC, ligand depletion binding, SEC

| Construct | promotor/transactivator | terminator | linker | C-tag | N-Tag | plasmid | resistance/selection marker | used in figure |
|---|---|---|---|---|---|---|---|---|
| AtLORE-GFP | CaMV35S promotor | CaMV35S terminator | 11x Glycin | eGFP | - | pGGPXB (based on pCB302, Ranf et al., 2015) | Kanamycin, BASTA | 1A, 1B; 3B, 4A, 5A, 6C, 6D, 7A, 7B, S2 |
| AtLORE-Km-GFP | CaMV35S promotor | CaMV35S terminator | 11x Glycin | eGFP | - | pGGPXB (based on pCB302, Ranf et al., 2015) | Kanamycin, BASTA | 1A, 1B, 2B, 3B, 5A, 6D, 7A, 7B, S2 |
| AtLORE-mCherry | CaMV35S promotor | CaMV35S terminator | 11x Glycin | mCherry | - | pGGPXB (based on pCB302, Ranf et al., 2015) | Kanamycin, BASTA | 1B |
| AtLORE-Km-mCherry | CaMV35S promotor | CaMV35S terminator | 11x Glycin | mCherry | - | pGGPXB (based on pCB302, Ranf et al., 2015) | Kanamycin, BASTA | 1B |
| AtLORE-HA | CaMV35S promotor | CaMV35S terminator | 6x Glycin | Hemagglutinin (HA) | - | pGGPXB (based on pCB302, Ranf et al., 2015) | Kanamycin, BASTA | 7C, 7D, S10 |
| AtLORE-Km-HA | CaMV35S promotor | CaMV35S terminator | 6x Glycin | Hemagglutinin (HA) | - | pGGPXB (based on pCB302, Ranf et al., 2015) | Kanamycin, BASTA | 2B, 7C, 7D, S10 |
| cytosolic mCherry | CaMV35S promotor | CaMV35S terminator | - | mCherry | - | pGGPXB (based on pCB302, Ranf et al., 2015) | Kanamycin, BASTA | 1B, 7A, 7B |
| cytosolic GFP | CaMV35S promotor | CaMV35S terminator | - | eGFP | - | pGGPXB (based on pCB302, Ranf et al., 2015) | Kanamycin, BASTA | 2B |
| AtLORE-ECD-TMD-GFP | CaMV35S promotor | CaMV35S terminator | 11x Glycin | eGFP | - | pGGPXB (based on pCB302, Ranf et al., 2015) | Kanamycin, BASTA | 2B |
| AtLORE-ECD-GFP | CaMV35S promotor | CaMV35S terminator | 11x Glycin | eGFP | - | pGGPXB (based on pCB302, Ranf et al., 2015) | Kanamycin, BASTA | 2B |
| AtLORE-ICD-Km-GFP | CaMV35S promotor | CaMV35S terminator | 11x Glycin | eGFP | - | pGGPXB (based on pCB302, Ranf et al., 2015) | Kanamycin, BASTA | 2B |
| AtLORE-TMD-ICD-Km-GFP | CaMV35S promotor | CaMV35S terminator | 11x Glycin | eGFP | - | pGGPXB (based on pCB302, Ranf et al., 2015) | Kanamycin, BASTA | 2B |
| AlyrLORE-GFP | CaMV35S promotor | CaMV35S terminator | 11x Glycin | eGFP | - | pGGPXB (based on pCB302, Ranf et al., 2015) | Kanamycin, BASTA | 3B,4A,5A, 6C, 6D |
| AhalLORE-GFP | CaMV35S promotor | CaMV35S terminator | 11x Glycin | eGFP | - | pGGPXB (based on pCB302, Ranf et al., 2015) | Kanamycin, BASTA | 3B,5A |
| CrubLORE-GFP | CaMV35S promotor | CaMV35S terminator | 11x Glycin | eGFP | - | pGGPXB (based on pCB302, Ranf et al., 2015) | Kanamycin, BASTA | 3B,5A |
| AtLORE-DSAlyrICD-GFP | CaMV35S promotor | CaMV35S terminator | 11x Glycin | eGFP | - | pGGPXB (based on pCB302, Ranf et al., 2015) | Kanamycin, BASTA | 4A, 6D |
| AtLORE-DSAlyrECD-TMD-GFP | CaMV35S promotor | CaMV35S terminator | 11x Glycin | eGFP | - | pGGPXB (based on pCB302, Ranf et al., 2015) | Kanamycin, BASTA | 4A, 6D |
| AtLORE-ECD-mCherry | CaMV35S promotor | CaMV35S terminator | 11x Glycin | mCherry | - | pGGPXB (based on pCB302, Ranf et al., 2015) | Kanamycin, BASTA | 4C, S7A |
| CrubLORE-ECD-mCherry | CaMV35S promotor | CaMV35S terminator | 11x Glycin | mCherry | - | pGGPXB (based on pCB302, Ranf et al., 2015) | Kanamycin, BASTA | 4C, S7A |
| AlyrLORE-ECD-mCherry | CaMV35S promotor | CaMV35S terminator | 11x Glycin | mCherry | - | pGGPXB (based on pCB302, Ranf et al., 2015) | Kanamycin, BASTA | 4C, S7A |
| AhalLORE-ECD-mCherry | CaMV35S promotor | CaMV35S terminator | 11x Glycin | mCherry | - | pGGPXB (based on pCB302, Ranf et al., 2015) | Kanamycin, BASTA | 4C, S7A |
| AtSD1-23i-ECD-mCherry | CaMV35S promotor | CaMV35S terminator | 11x Glycin | mCherry | - | pGGPXB (based on pCB302, Ranf et al., 2015) | Kanamycin, BASTA | 4C, S7A |
| apoplastic mCherry | CaMV35S promotor | CaMV35S terminator | - | mCherry | - | pGGPXB (based on pCB302, Ranf et al., 2015) | Kanamycin, BASTA | 4C, S7A |
| TwinStrep-TEV-AtLORE-ECD-HA | CaMV35S promotor | CaMV35S terminator | GS linker (GGSGGSG) | Hemagglutinin (HA) | TwinStrep-5xGly-TEV | pGGPXhc (based on pCB302, Ranf et al., 2015) | Kanamycin | 4D, S7C, S7D |
| TwinStrep-TEV-AlyrLORE-ECD-HA | CaMV35S promotor | CaMV35S terminator | GS linker (GGSGGSG) | Hemagglutinin (HA) | TwinStrep-5xGly-TEV | pGGPXhc (based on pCB302, Ranf et al., 2015) | Kanamycin | 4D, S7C, S7D |
| AtLORE-DSAlyrLL-GFP | CaMV35S promotor | CaMV35S terminator | 11x Glycin | eGFP | - | pGGPXB (based on pCB302, Ranf et al., 2015) | Kanamycin, BASTA | 6C, 6D |
| AtLORE-DSAlyrEP-GFP | CaMV35S promotor | CaMV35S terminator | 11x Glycin | eGFP | - | pGGPXB (based on pCB302, Ranf et al., 2015) | Kanamycin, BASTA | 6C, 6D |
| AtLORE-ECD-TMD-HA | CaMV35S promotor | CaMV35S terminator | 6x Glycin | Hemagglutinin (HA) | - | pGGPXB (based on pCB302, Ranf et al., 2015) | Kanamycin, BASTA | 7A, 7C, 7D, S10 |
| AtLORE-TMD-ICD-HA | CaMV35S promotor | CaMV35S terminator | 6x Glycin | Hemagglutinin (HA) | - | pGGPXB (based on pCB302, Ranf et al., 2015) | Kanamycin, BASTA | 7B |
| AlyrLORE-ECD-TMD-HA | CaMV35S promotor | CaMV35S terminator | 6x Glycin | Hemagglutinin (HA) | - | pGGPXB (based on pCB302, Ranf et al., 2015) | Kanamycin, BASTA | 7C, 7D, S10 |
| HA-AtLORE-ECD | CaMV35S promotor | CaMV35S terminator | 5x Glycin | - | Hemagglutinin (HA) | pGGPXB (based on pCB302, Ranf et al., 2015) | Kanamycin, BASTA | 7C, 7D, S10 |
| AtLORE-Km-SPYCE | CaMV35S promotor | CaMV35S terminator | 5x Glycin | SPYCE (c-myc -eYFP$^{N(aa1-84)}$) | - | pGGPXB (based on pCB302, Ranf et al., 2015) | Kanamycin, BASTA | S3A, S3B, S3C |
| AtLORE-Km-SPYNE | CaMV35S promotor | CaMV35S terminator | 5x Glycin | SPYNE (HA-eYFP$^{C(aa85-257)}$) | - | pGGPXB (based on pCB302, Ranf et al., 2015) | Kanamycin, BASTA | S3A, S3B, S3C |
| cytosolic SPYCE | CaMV35S promotor | CaMV35S terminator | - | SPYCE (c-myc -eYFP$^{N(aa1-84)}$) | - | pGGPXB (based on pCB302, Ranf et al., 2015) | Kanamycin, BASTA | S3A, S3B, S3C |

For FLIM

| Construct | promotor/transactivator | terminator | linker | C-tag | N-Tag | plasmid | resistance/selection marker | used in figure |
|---|---|---|---|---|---|---|---|---|
| AtLORE-Km-GFP | XVE (estradiol inducible transactivator casette) | T3A terminator | attB2 site (AADPAFLYKVVDHSLIG) | GFP | - | pABindGFP (Bleckmann et al., 2009) | Spectinomycin, Hygromycin | 1C, 1D, 2C, 6B, S3D, S5B, S8B, S8C |
| AtLORE-Km-mCherry | XVE (estradiol inducible transactivator casette) | T3A terminator | attB2 site (AADPAFLYKVVDHSLIG) | mCherry | - | pABindmCherry (Bleckmann et al., 2009) | Spectinomycin, Hygromycin | 1C, 1D, 2C, 5B, 6B, S3D, S8B |
| AtLORE-Km-GFP-mCherry | XVE (estradiol inducible transactivator casette) | T3A terminator | attB2 site (AADPAFLYKVVDHSLIG) | GFP-mCherry | - | pABindFRET (Bleckmann et al., 2009) | Spectinomycin, Hygromycin | 1C, 1D, 2C, 6B, S3D, S8B, S8C |
| AtGST-mCherry | XVE (estradiol inducible transactivator casette) | T3A terminator | attB2 site (AADPAFLYKVVDHSLIG) | mCherry | - | pABindmCherry (Bleckmann et al., 2009) | Spectinomycin, Hygromycin | 1C, 5B, 5C, 5D, 6B, S3D, S8B, S8C |
| AtLORE-ECD-TMD-GFP | XVE (estradiol inducible transactivator casette) | T3A terminator | attB2 site (AADPAFLYKVVDHSLIG) | GFP | - | pABindGFP (Bleckmann et al., 2009) | Spectinomycin, Hygromycin | 2D |
| AtLORE-ECD-TMD-mCherry | XVE (estradiol inducible transactivator casette) | T3A terminator | attB2 site (AADPAFLYKVVDHSLIG) | mCherry | - | pABindmCherry (Bleckmann et al., 2009) | Spectinomycin, Hygromycin | 2C, 2D, S4 |
| AtLORE-TMD-ICD-mCherry | XVE (estradiol inducible transactivator casette) | T3A terminator | attB2 site (AADPAFLYKVVDHSLIG) | mCherry | - | pABindmCherry (Bleckmann et al., 2009) | Spectinomycin, Hygromycin | 2C, 2D, S4 |
| AlyrLORE-Km-GFP | XVE (estradiol inducible transactivator casette) | T3A terminator | attB2 site (AADPAFLYKVVDHSLIG) | GFP | - | pABindGFP (Bleckmann et al., 2009) | Spectinomycin, Hygromycin | 5B, 6B, S5B, S8B, S8C |
| AlyrLORE-Km-mCherry | XVE (estradiol inducible transactivator casette) | T3A terminator | attB2 site (AADPAFLYKVVDHSLIG) | mCherry | - | pABindmCherry (Bleckmann et al., 2009) | Spectinomycin, Hygromycin | 5B, 6B, S8B |
| AlyrLORE-Km-GFP-mCherry | XVE (estradiol inducible transactivator casette) | T3A terminator | attB2 site (AADPAFLYKVVDHSLIG) | GFP-mCherry | - | pABindFRET (Bleckmann et al., 2009) | Spectinomycin, Hygromycin | 5B |
| AhalLOREi-Km-GFP | XVE (estradiol inducible transactivator casette) | T3A terminator | attB2 site (AADPAFLYKVVDHSLIG) | GFP | - | pABindGFP (Bleckmann et al., 2009) | Spectinomycin, Hygromycin | 5C, S5B |
| AhalLOREi-Km-mCherry | XVE (estradiol inducible transactivator casette) | T3A terminator | attB2 site (AADPAFLYKVVDHSLIG) | mCherry | - | pABindmCherry (Bleckmann et al., 2009) | Spectinomycin, Hygromycin | 5C |
| AhalLOREi-Km-GFP-mCherry | XVE (estradiol inducible transactivator casette) | T3A terminator | attB2 site (AADPAFLYKVVDHSLIG) | GFP-mCherry | - | pABindFRET (Bleckmann et al., 2009) | Spectinomycin, Hygromycin | 5C |
| CrubLORE-Km-GFP | XVE (estradiol inducible transactivator casette) | T3A terminator | attB2 site (AADPAFLYKVVDHSLIG) | GFP | - | pABindGFP (Bleckmann et al., 2009) | Spectinomycin, Hygromycin | 5D, S5B |
| CrubLORE-Km-mCherry | XVE (estradiol inducible transactivator casette) | T3A terminator | attB2 site (AADPAFLYKVVDHSLIG) | mCherry | - | pABindmCherry (Bleckmann et al., 2009) | Spectinomycin, Hygromycin | 5D |
| CrubLORE-Km-GFP-mCherry | XVE (estradiol inducible transactivator casette) | T3A terminator | attB2 site (AADPAFLYKVVDHSLIG) | GFP-mCherry | - | pABindFRET (Bleckmann et al., 2009) | Spectinomycin, Hygromycin | 5D |
| AtLORE-DSAlyrECD-TMD-Km-GFP | XVE (estradiol inducible transactivator casette) | T3A terminator | attB2 site (AADPAFLYKVVDHSLIG) | GFP | - | pABindGFP (Bleckmann et al., 2009) | Spectinomycin, Hygromycin | 6B, S9 |
| AtLORE-DSAlyrECD-TMD-Km-mCherry | XVE (estradiol inducible transactivator casette) | T3A terminator | attB2 site (AADPAFLYKVVDHSLIG) | mCherry | - | pABindmCherry (Bleckmann et al., 2009) | Spectinomycin, Hygromycin | 6B |
| AtLORE-DSAlyrECD-Km-GFP | XVE (estradiol inducible transactivator casette) | T3A terminator | attB2 site (AADPAFLYKVVDHSLIG) | GFP | - | pABindGFP (Bleckmann et al., 2009) | Spectinomycin, Hygromycin | 6B, S9 |
| AtLORE-DSAlyrECD-Km-mCherry | XVE (estradiol inducible transactivator casette) | T3A terminator | attB2 site (AADPAFLYKVVDHSLIG) | mCherry | - | pABindmCherry (Bleckmann et al., 2009) | Spectinomycin, Hygromycin | 6B |
| AtLORE-DSAlyrLL-Km-GFP | XVE (estradiol inducible transactivator casette) | T3A terminator | attB2 site (AADPAFLYKVVDHSLIG) | GFP | - | pABindGFP (Bleckmann et al., 2009) | Spectinomycin, Hygromycin | 6B,S9 |
| AtLORE-DSAlyrLL-Km-mCherry | XVE (estradiol inducible transactivator casette) | T3A terminator | attB2 site (AADPAFLYKVVDHSLIG) | mCherry | - | pABindmCherry (Bleckmann et al., 2009) | Spectinomycin, Hygromycin | 6B, S8C |
| AtLORE-DSAlyrEP-Km-GFP | XVE (estradiol inducible transactivator casette) | T3A terminator | attB2 site (AADPAFLYKVVDHSLIG) | GFP | - | pABindGFP (Bleckmann et al., 2009) | Spectinomycin, Hygromycin | 6B, S8C, S9 |
| AtLORE-DSAlyrEP-Km-mCherry | XVE (estradiol inducible transactivator casette) | T3A terminator | attB2 site (AADPAFLYKVVDHSLIG) | mCherry | - | pABindmCherry (Bleckmann et al., 2009) | Spectinomycin, Hygromycin | 6B |
| AtLORE-DSAlyrTMD-ICD-Km-GFP | XVE (estradiol inducible transactivator casette) | T3A terminator | attB2 site (AADPAFLYKVVDHSLIG) | GFP | - | pABindGFP (Bleckmann et al., 2009) | Spectinomycin, Hygromycin | S8C |
| AtLORE-DSAlyrTMD-ICD-Km-mCherry | XVE (estradiol inducible transactivator casette) | T3A terminator | attB2 site (AADPAFLYKVVDHSLIG) | mCherry | - | pABindmCherry (Bleckmann et al., 2009) | Spectinomycin, Hygromycin | S8C |
| AtLORE-DSAlyrL1-Km-GFP | XVE (estradiol inducible transactivator casette) | T3A terminator | attB2 site (AADPAFLYKVVDHSLIG) | GFP | - | pABindGFP (Bleckmann et al., 2009) | Spectinomycin, Hygromycin | S8B, S9 |
| AtLORE-DSAlyrL1-Km-mCherry | XVE (estradiol inducible transactivator casette) | T3A terminator | attB2 site (AADPAFLYKVVDHSLIG) | mCherry | - | pABindmCherry (Bleckmann et al., 2009) | Spectinomycin, Hygromycin | S8B |
| AtLORE-DSAlyrL2-Km-GFP | XVE (estradiol inducible transactivator casette) | T3A terminator | attB2 site (AADPAFLYKVVDHSLIG) | GFP | - | pABindGFP (Bleckmann et al., 2009) | Spectinomycin, Hygromycin | S8B, S9 |
| AtLORE-DSAlyrL2-Km-mCherry | XVE (estradiol inducible transactivator casette) | T3A terminator | attB2 site (AADPAFLYKVVDHSLIG) | mCherry | - | pABindmCherry (Bleckmann et al., 2009) | Spectinomycin, Hygromycin | S8B |
| AtLORE-DSAlyrE-Km-GFP | XVE (estradiol inducible transactivator casette) | T3A terminator | attB2 site (AADPAFLYKVVDHSLIG) | GFP | - | pABindGFP (Bleckmann et al., 2009) | Spectinomycin, Hygromycin | S8B, S9 |
| AtLORE-DSAlyrE-Km-mCherry | XVE (estradiol inducible transactivator casette) | T3A terminator | attB2 site (AADPAFLYKVVDHSLIG) | mCherry | - | pABindmCherry (Bleckmann et al., 2009) | Spectinomycin, Hygromycin | S8B |
| AtLORE-DSAlyrP-Km-GFP | XVE (estradiol inducible transactivator casette) | T3A terminator | attB2 site (AADPAFLYKVVDHSLIG) | GFP | - | pABindGFP (Bleckmann et al., 2009) | Spectinomycin, Hygromycin | S8B, S9 |
| AtLORE-DSAlyrP-Km-mCherry | XVE (estradiol inducible transactivator casette) | T3A terminator | attB2 site (AADPAFLYKVVDHSLIG) | mCherry | - | pABindmCherry (Bleckmann et al., 2009) | Spectinomycin, Hygromycin | S8B |

For stable transgenic lines

| Construct | promotor/transactivator | terminator | linker | C-tag | N-Tag | plasmid | resistance/selection marker | used in figure |
|---|---|---|---|---|---|---|---|---|
| AtLORE | pAtLORE (Ranf *et al* ., 2015) | tHSP (de Felippes *et al* ., 2020) | - | - | - | pGGPXhcFR (based on pCB302, Ranf et al., 2015) | Kanamycin, FastRed | 3C, 3D |
| AlyrLORE | pAtLORE (Ranf *et al* ., 2015) | tHSP (de Felippes *et al* ., 2020) | - | - | - | pGGPXhcFR (based on pCB302, Ranf et al., 2015) | Kanamycin, FastRed | 3C, 3D |
| CrubLORE | pAtLORE (Ranf *et al* ., 2015) | tHSP (de Felippes *et al* ., 2020) | - | - | - | pGGPXhcFR (based on pCB302, Ranf et al., 2015) | Kanamycin, FastRed | 3C, 3D |
| AtLORE-ECD-TMD | CaMV35S promotor | CaMV35S terminator | 11x Glycin | mCherry | - | pGGPXhcFR (based on pCB302, Ranf et al., 2015) | Kanamycin, FastRed | 7E, S11 |
| AtLORE-ECD | CaMV35S promotor | CaMV35S terminator | 11x Glycin | mCherry | - | pGGPXhcFR (based on pCB302, Ranf et al., 2015) | Kanamycin, FastRed | 7E, S11 |

**Table S5 DNA and protein sequence sources**

| Gene | Locus ID/Gene model | Source gene information | Protein ID | Source protein information | Uniprot ID |
|---|---|---|---|---|---|
| **LORE orthologs used in this study** | | | | | |
| *AtLORE* | AT1G61380 | TAIR (https://www.arabidopsis.org/servlets/TairObject?id=136677&type=locus) | AT1G61380.1 | TAIR (https://www.arabidopsis.org/servlets/TairObject?id=1009109351&type=aa_sequence) | O64782 |
| *AhalLORE* | Araha.6790s0007.1 | Phytozome (https://phytozome-next.jgi.doe.gov/report/transcript/Ahalleri_v1_1/Araha.6790s0007.1) | Araha.6790s0007.1.p | See Source Gene information | no entry |
| *AlyrLORE* | AL2G14950, ARALYDRAFT_475185 (or 9322615) | Phytozome (https://phytozome-next.jgi.doe.gov/report/gene/Alyrata_v2_1/AL2G14950)<br>NCBI (https://www.ncbi.nlm.nih.gov/gene/?term=ARALYDRAFT_475185) | XP_020890046.1 | NCBI (https://www.ncbi.nlm.nih.gov/protein/XP_020890046.1) | D7KW16 |
| *CrubLORE* | CARUB_v10021901mg (or 17894778) | NCBI (https://www.ncbi.nlm.nih.gov/gene/?term=CARUB_v10021901mg) | XP_006301473 | NCBI (https://www.ncbi.nlm.nih.gov/protein/XP_006301473.1) | R0GFC4 |
| **Closest LORE paralogs in *A. thaliana*** | | | | | |
| *AtSD1-23* | AT1G61390 | TAIR (https://www.arabidopsis.org/servlets/TairObject?id=136672&type=locus) | AT1G61390.1 | TAIR (https://www.arabidopsis.org/servlets/TairObject?type=aa_sequence&id=1009109352) | O64781 |
| *AtSD1-27* | AT1G61370.1 | TAIR (https://www.arabidopsis.org/servlets/TairObject?id=136678&type=locus) | AT1G61370.1 | TAIR (https://www.arabidopsis.org/servlets/TairObject?type=aa_sequence&id=1009109354) | O64783 |
| *AtSD1-30* | AT1G61360.1 | TAIR (https://www.arabidopsis.org/servlets/TairObject?id=136661&type=locus) | AT1G61360.1 | TAIR (https://www.arabidopsis.org/servlets/TairObject?type=aa_sequence&id=1009109356) | O64784 |
| **Well-studied SRKs from *B. rapa* (Ma et al., 2016; Murase et al.,2020** | | | | | |
| *BraSRK8* | AB257127 | NCBI (https://www.ncbi.nlm.nih.gov/nuccore/AB257127.1) | BAF91375 | NCBI (https://www.ncbi.nlm.nih.gov/protein/BAF91375) | A8QWG5 |
| *BraSKR9* | D88193 | NCBI (https://www.ncbi.nlm.nih.gov/nuccore/D88193) | BAA21132.1 | NCBI (https://www.ncbi.nlm.nih.gov/protein/BAA21132.1) | O23745 |
| **Other PRR-RLKs and RLCKs** | | | | | |
| *AtFLS2* | AT5G46330.1 | TAIR (https://www.arabidopsis.org/servlets/TairObject?id=134136&type=locus) | AT5G46330.1 | TAIR (https://www.arabidopsis.org/servlets/TairObject?type=aa_sequence&id=1009133864) | Q9FL28 |
| *AtBAK1* | AT4G33430.2 | TAIR (https://www.arabidopsis.org/servlets/TairObject?id=127207&type=locus) | AT4G33430.2 | TAIR (https://www.arabidopsis.org/servlets/TairObject?type=aa_sequence&id=6530309380) | Q94F62 |
| *AtEFR* | AT5G20480.1 | TAIR (https://www.arabidopsis.org/servlets/TairObject?id=131371&type=locus) | AT5G20480.1 | TAIR (https://www.arabidopsis.org/servlets/TairObject?type=aa_sequence&id=1009134859) | C0LGT6 |
| *AtCERK1* | AT3G21630.1 | TAIR (https://www.arabidopsis.org/servlets/TairObject?id=38332&type=locus) | AT3G21630.1 | TAIR (https://www.arabidopsis.org/servlets/TairObject?type=aa_sequence&id=1009119534) | A8R7E6 |
| *AtBIK1* | AT2G39660.1 | TAIR (https://www.arabidopsis.org/servlets/TairObject?id=31443&type=locus) | AT2G39660.1 | TAIR (https://www.arabidopsis.org/servlets/TairObject?id=31443&type=locus) | O48814 |

**Top hits BLAST of AtLORE and AtSD1-23 against *A.lyrata, A. halleri, C.rubella* and *B. rapa* genomes (Phytozome)**

| Locus ID/Gene model | Source gene information |
|---|---|
| Brara.I01503.1 | Phytozome (https://phytozome-next.jgi.doe.gov/) |
| Brara.A02537.1 | Phytozome (https://phytozome-next.jgi.doe.gov/) |
| Brara.I01502.1 | Phytozome (https://phytozome-next.jgi.doe.gov/) |
| Carub.0002s0379.1 | Phytozome (https://phytozome-next.jgi.doe.gov/) |
| AL2G14830.t1 | Phytozome (https://phytozome-next.jgi.doe.gov/) |
| Ah2G04790.2 | Phytozome (https://phytozome-next.jgi.doe.gov/) |

**SD-RLKs (or G-type lectin RLK) from tomato and rice with similar domain achitecture than *At* LORE (Lectin, EGF, PAN domain) as annotated by Teixeira *et al.,* 2018**

| Locus ID/Gene model | Source gene information |
|---|---|
| Solyc02g079640.2.1 | iTAK (http://itak.feilab.net/cgi-bin/itak/index.cgi) |
| Solyc04g058110.2.1 | iTAK (http://itak.feilab.net/cgi-bin/itak/index.cgi) |
| Solyc07g063770.2.1 | iTAK (http://itak.feilab.net/cgi-bin/itak/index.cgi) |
| Solyc11g005630.1.1 | iTAK (http://itak.feilab.net/cgi-bin/itak/index.cgi) |
| Solyc10g006710.2.1 | iTAK (http://itak.feilab.net/cgi-bin/itak/index.cgi) |
| LOC_Os01g47810.1 | iTAK (http://itak.feilab.net/cgi-bin/itak/index.cgi) |
| LOC_Os01g47840.1 | iTAK (http://itak.feilab.net/cgi-bin/itak/index.cgi) |
| LOC_Os01g57560.1 | iTAK (http://itak.feilab.net/cgi-bin/itak/index.cgi) |
| LOC_Os03g12150.1 | iTAK (http://itak.feilab.net/cgi-bin/itak/index.cgi) |
| LOC_Os03g30890.1 | iTAK (http://itak.feilab.net/cgi-bin/itak/index.cgi) |

**Table 6 Antibodies**

| Primary Antibody | | Secondary Antibody | | |
|---|---|---|---|---|
| Name | dilution | Name | Dilution | Used in Figure |
| anti-GFP 3H9 (ChromoTek) | 1:1000 | anti-rat-HRP (A9542, Sigma-Aldrich) | 1:20000 | Fig 1B, 2B |
| anti-mCherry 5F8 (ChromoTek) | 1:2000 | anti-rat-HRP (A9542, Sigma-Aldrich) | 1:20000 | Fig 1B |
| anti-HA 7C9 (ChromoTek) | 1:2000 | anti-rat-HRP (A9542, Sigma-Aldrich) | 1:20000 | Fig S3B, S7, S10 |
| c-Myc 9E10 (Santa Cruz) | 1:500 | anti-mouse sc-2031 (Santa Cruz) | 1:5000 | Fig S3B |
| anti-HA-HRP 3F10 (Sigma-Aldrich) | 1:2000 | none | | Fig 2B |
| StrepMAB-classic (IBA Lifesciences) | 1:10000 | anti-mouse-HRP (A9044, Sigma-Aldrich) | 1:20000 | Fig S7C |
